# Bridging the gap between genes and language deficits in schizophrenia: an oscillopathic approach

**DOI:** 10.1101/043547

**Authors:** Elliot Murphy, Antonio Benítez-Burraco

## Abstract

Schizophrenia is characterised by marked language deficits, but it is not clear how these deficits arise from gene mutations linked to or associated with the disease. The goal of this paper is to aid the bridging of the gap between genes and schizophrenia and, ultimately, give support to the view that it represents an abnormal ontogenetic itinerary for the human faculty of language, heavily rooted in the evolutionary processes that brought about modern language. To that end we will focus on how the schizophrenia brain processes language and, particularly, on its distinctive oscillatory profile during language processing: We will argue that brain rhythms constitute the best route to interpret language deficits in this condition and map them to neural dysfunction and risk alleles of the genes. Additionally, we will show that candidate genes for schizophrenia are overrepresented among the set of genes that are believed are important for the evolution of human language. These genes crucially include (and are related to) genes involved in brain rhythmicity. We will claim that this translational effort and the links we uncover may help develop an understanding of language evolution, along with the aetiology of schizophrenia, its clinical/linguistic profile, and its high prevalence among modern populations.

## 1. Introduction

Schizophrenia is a pervasive neurodevelopmental disorder entailing several (and severe) social and cognitive deficits (van Os and Kapur 2009). Usually, people with schizophrenia exhibit language problems at all levels, from phonology to pragmatics, which coalesce into problems for speech perception (auditory verbal hallucinations), abnormal speech production (formal thought disorder), and production of abnormal linguistic content (delusions) (Figure 1), which are the hallmarks of the disease in the domain of language (Stephane et al. 2007, Stephane et al. 2014). In turn, language dysfunction has been hypothesised to result from the impairment of some basic process (or processes), either linguistic or non-linguistic (e.g. semantic memory and/or working memory and executive function) (Kuperberg, 2010). Importantly, although schizophrenia is commonly defined as a disturbance of thought or selfhood, some authors claim that most of its distinctive symptoms may arise from language dysfunction; in particular, from failures in language-mediated forms of meaning (Hinzen and Rosselló 2015). These perspectives crucially depart from Bleuler’s (1911) original definition of schizophrenia which kept the cognitive and speech-related symptoms separated.

**Figure 1.**
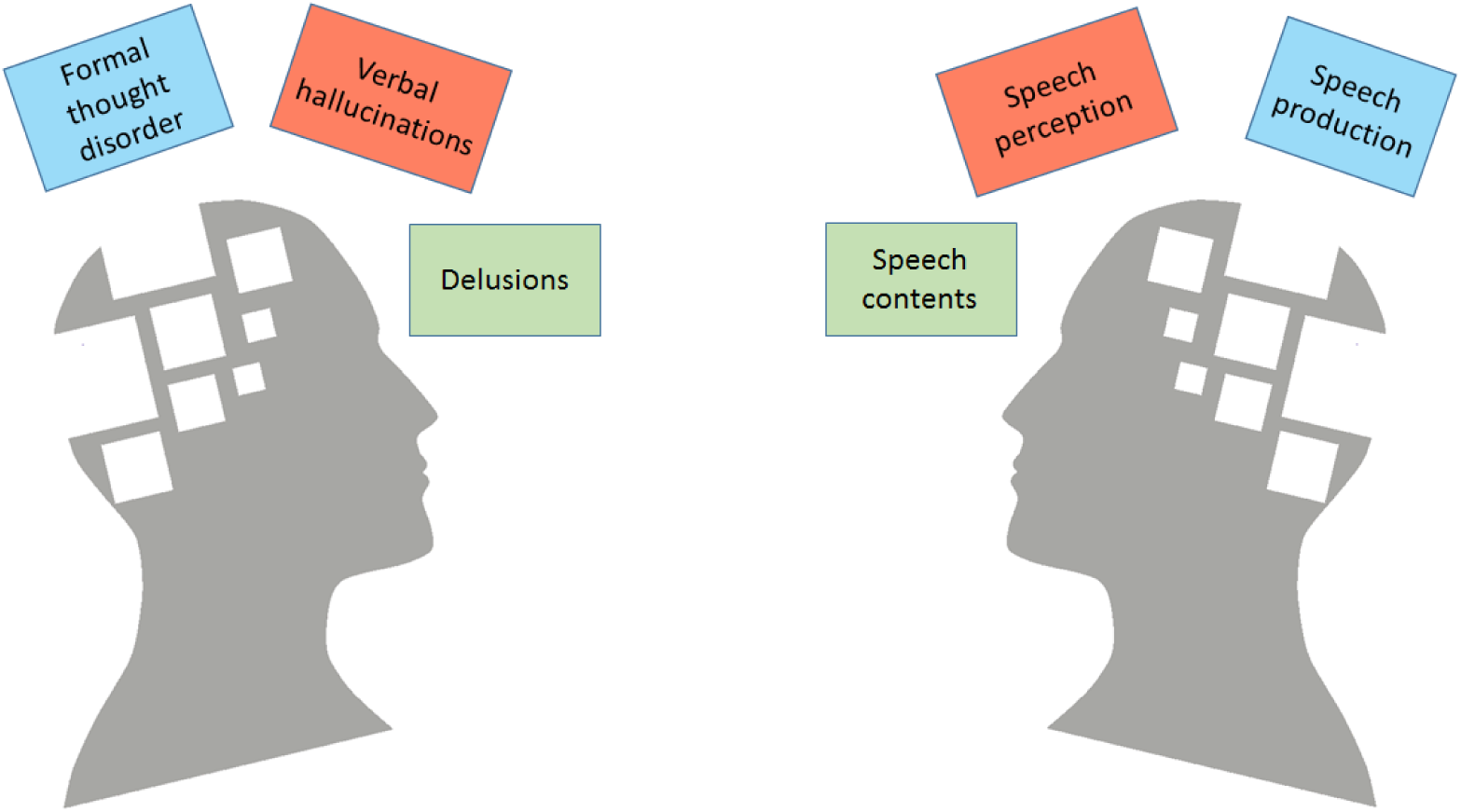
The three main positive symptoms of schizophrenia derived as a disruption in normal linguistic cognition (adapted from Hinzen and Rosselló 2015; the scheme of the head was taken from http://journal.frontiersin.org/journal/psychiatry/section/schizophrenia).

Schizophrenia involves atypical brain development and wiring during growth, which results in a distinctive neurocognitive profile. Typically, changes in ventricle size, gray matter density, whole-brain volumes, and interconnection patterns are observed, seemingly resulting from altered neuroplasticity and vulnerability of inhibitory cortical circuits involving interneurons (Bakhshi and Chance 2015) and, in particular, from disruption in brain connectivity driven by a reduction in dendritic spines on cortical pyramidal neurons (Cannon 2015). Not surprisingly, structural and functional abnormalities in brain regions involved in language processing are observed in schizophrenics: typically, left hemisphere dominance of language processing and left-right communication are disturbed in them (Li et al. 2009, Li et al. 2012).

At the same time, there is ample evidence that schizophrenia is caused by a complex interaction between genetic, epigenetic and environmental factors. To date, schizophrenia has been linked or associated to mutations in an extensive number of genes (see O’Tuathaigh et al. 2012, Flint and Munafò 2014, and McCarthy et al. 2014 for recent reviews). Many of them point to specific pathways (like glutamatergic, GABAergic and cholinergic pathways, the neuregulin signalling pathway, and the Akt/GSK-3 pathway) and to specific neural mechanisms (like those involving dendritic spines and synaptic terminals, synapses, gray matter development, and neural plasticity) (Buonanno 2010, Karam et al. 2010, Bennet et al. 2011, Hall et al. 2015). However, the gap between genes, brain abnormalities, and cognitive dysfunction in schizophrenia still remains open, particularly regarding its distinctive linguistic profile.

The goal of this paper is to contribute to the bridging of this gap between genes and schizophrenia. To that end we will focus on how the schizophrenic brain processes language and, more specifically, on its distinctive oscillatory profile during language processing. We believe that brain rhythms may constitute the best route to interpreting language deficits in schizophrenia and mapping them to neural dysfunction and risk alleles of the genes. First, because brain rhythms are connected to some computational primitives of language (see Murphy 2015a for discussion), they allow to explain, and not just to describe, how the brain processes language. Consequently, they are not faced with the sort of problems that most neurolinguistics research – heavily focused on the maps provided by structural and functional neuroimagining – has to overcome; namely, a granularity problem (that is, neuroscientific studies of language operate at different scales from linguistic analyses) and an incommensurability problem (that is, the basic components of linguistic theory cannot be reduced or paired up with the basic biological units identified by neuroscience) (see Poeppel and Embick 2005 for discussion). Second, cognitive disorders can be conceived of as oscillopathies, or pathological variations of the normal profile of brain rhythmicity (Buzsáki et al. 2013). Current understanding suggests that schizophrenia is characterized by asynchronous neural oscillations, and particularly, by an inhibitory interneuron dysfunction (Moran and Hong 2011, Pittman-Polleta et al. 2015). Third, brain rhythms are heritable components of brain function (Linkenkaer-Hansen et al. 2007), also in pathological conditions (see Hall et al, 2011 for schizophrenia). Fourth, brain rhythms connect to both aspects of human biology conserved across species and aspects of human biology known to vary within the species, allowing for an evolutionary-developmental (evo-devo) approach to human cognition, including language abilities (Benítez-Burraco and Boeckx 2014). Accordingly, the hierarchy of brain oscillations has remained remarkably preserved within mammals during their evolution (Buzsáki et al. 2013), to the extent that the human pattern of brain activity can be conceived of as a slight variation of the patterns observed in other primates. Similarly, different cognitive disorders have proven to correlate with distinct, disorder-specific patterns of anomalous brain activity (Buzsáki and Watson 2012).

If we are on the right track, we expect that examining the genes related to brain dysrhythmias in schizophrenia and interpreting language deficits in this condition as oscillopathic features will allow the construction of successful endophenotypes of schizophrenia, and ultimately help achieve a better treatment of those affected. Moreover, we will advocate an evo-devo approach to schizophrenia that should allow for a better understanding of its nature, origins, and prevalence among human populations. More generally, we believe that this translational effort should also contribute to the growing understanding of the human faculty of language, set against an evolving dynamic model of mental computation, and of language evolution, in turn set against a reorganizational model of the evolution of cognition. Concerning the neurobiological foundations of language, we expect our approach will cast light on how the brain processes language. In doing so, our attempts of translating language into a grammar of brain rhythms will heavily rely on the model developed in Murphy (2015a), where it is argued that brain rhythms are the suitable neuronal processes which can capture the computational properties of the human language faculty and advance a new model of linguistic computation. Regarding the evolution of the human language, we also expect to shed some much-needed light on the changes that brought about our distinctive mode of cognition, known to be impaired in schizophrenics. In doing so, we will build on our ongoing research into the origins and development of the human faculty of language. According to our view, human language-readiness resulted from subtle changes in the developmental trajectory of the hominin brain/skull that were brought about by modifications in some of the involved genes. These changes seemingly modified the primate pattern of cortical inhibition and brought about the sort of human-specific pattern of long-distance connections across the brain that enable us to form and exploit cross-modular concepts (see Boeckx and Benítez-Burraco 2014a,b, Benítez-Burraco and Boeckx, 2015a for details), both of which are aspects that are targeted in schizophrenia (Morice and McNicol, 1985, Horn et al. 2012, Jiang et al. 2015).

The paper is structured as follows. First, we provide a general account of language deficits in schizophrenia and the attested anomalies in brain structure and function in schizophrenics in connection to language processing. Next we focus on brain rhythms and advance a tentative oscillopathic model of language deficits in schizophrenia. Then we move to the genes. We first review some of the candidates for schizophrenia that may help to explain its abnormal profile of brain rhythmicity. Afterwards, we examine the oscillopathic nature of language deficits in schizophrenia from a broader, evo-devo perspective. Accordingly, we will examine candidate genes for this condition under the lens of the genetic changes known to have occurred after our split from extinct hominins. The last section of the paper provides a summary of the topics discussed in previous sections. We will claim that delving into the oscillopathic nature of language deficits in schizophrenia will help us understand its distinctive neurocognitive profile, its origins, and its prevalence among modern populations, ultimately yielding enhanced insights into the nature of language and how our language-readiness evolved.

## 2. From language deficits to the brain in schizophrenia

Schizophrenics have been known to have disordered speech (McKenna and Oh 2005), but the most severe linguistic changes occur at the internal, conceptual level, where studies frequently examine patients who experience thoughts being ‘inserted’ into them from outside sources or ‘broadcast’ out of their minds and into other people’s (Crow 1980, Frith 1992). Accordingly, one of the most fundamental deficits seen in schizophrenic speech is ‘the initiating and accessing of new ideas in a discourse’ (Frith 1992: 111). Patients also sometimes hear their thoughts ‘echoed’, or spoken aloud, and are also known to experience third-person and second-person auditory hallucinations, with an external voice either discussing them or commenting on their actions (Ramsden 2013: 234-265). Such hallucinations are experienced by over 70% of patients (Sartorius et al. 1986). Schizophrenic speech can be deviant without being incomprehensible, as in cases of impoverished and inadequate speech content, tangential speech, repetitive speech, repeated self-reference, and illogicality (Frith 1992). Patients often experience difficulty translating their thoughts into speech, but the difficulties appear to be largely at the discourse level, and are not usually syntactically ill-formed (Andreasen et al. 1985). Frith and Allen’s (1988) review observed ‘a failure to structure discourse at higher levels’. A cardinal feature of the schizophrenic linguistic profile, then, appears to be problems with externalisation and the ability to consider the listener’s knowledge when formulating speech, the latter a problem which amounts to a pragmatic impairment. Abnormalities can also be detected with syntax, however, and this is where we will focus most of our attention. Schizophrenic patients exhibit fewer relative clauses (as their discourse difficulties would predict), shorter utterances, and less clausal embedding (Thomas et al. 1987, Fraser et al. 1986). Importantly, this relative lack of clausal embedding implies that patients do not engage in thoughts about mental states or Theory of Mind (Morice and Ingram 1982, Morice and McNicol 1986). An fMRI study presenting patients with complex sentences revealed reduced activation in the right posterior temporal and left superior frontal cortex in schizophrenic patients relative to normal controls (Kircher et al. 2005).

‘Jargon aphasia’, which schizophrenic speech appears similar to, involves the production of incomprehensible speech – what Kraeplin (1913) may have had in mind when discussing the *“incoherence of thought”* in schizophrenics – and is associated with damage to the parietal-temporal junction and the arcuate fasciculus (McCarthy & Warrington 1990). Patients with lesions of the dorsolateral prefrontal cortex repeat themselves and use simple sentences, similar to schizophrenics (Kaczmarek 1987). In contrast to normal left-lateralization of activity in fronto-temporal regions during language processing, a wide range of schizophrenic patients exhibit bilateral and right-lateralized activity (Weiss et al. 2005, Diederen et al. 2010). Angrilli and colleagues (2009) have relatedly proposed that, judging by evoked potentials, certain features of schizophrenia appear to be (partly) a failure of phonological left hemispheric dominance, since the above deficit in lateralization is specific to phonological processing, being absent in semantic and word recognition tasks. Schizophrenics have also been known to have pedantic and overformal speech, as when Harrow and Quinlan (1985) asked a patient to respond to the proverb “Don’t judge a book by its cover” and were told “A façade of regal compliance bides an aetiology of ire”. The almost Joycean levels of disregard for the listener these kinds of stilted responses exhibit likely reflect an inability to succinctly formulate thoughts. Other studies have documented deficits in semantic memory (Tamlyn et al. 1992, Wang et al. 2011) which arise at the level of sentential, but not nominal, organisation.

The major positive symptoms of schizophrenia may amount to disturbances in linguistic computation. Delusions appear capable of being reduced to the unwilled production of abnormal internal language (i.e. disruptions at the sensorimotor-computational interface), ‘formal thought disorder’ may reduce to such abnormal production operating without feedback control, and auditory hallucinations seem remarkably close to speech perception disorders and Person confusions (see Hinzen and Rosselló 2015, and Figure 1). Indeed, formal thought disorder has been associated with gray matter volume reductions in language-related brain regions like Broca’s area and the superior temporal gyrus (Sans-Sansa et al. 2013), and seems capable of being captured in terms of a disrupted conceptual-computational system interface Schizophrenics additionally have a variety of problems with referential forms of propositional meaning, with deictic and definite noun phrases being impoverished relative to non-definite phrases (McKenna and Oh 2005). Since the deictic shifting and misperceptions of Personhood in schizophrenia appear to coexist with a narrowing and limiting of grammatical complexity, it may be that the neuronal operations which give rise to phrase structure building also generate these language-specific representations (see Murphy 2015b for discussion of how a number of grammatical and semantic structures co-vary). Finally, if delusions are false *beliefs*, and hallucinations are false *perceptions*, then we would predict that the neuronal dynamics observed for schizophrenics with one but not both of these symptoms would reflect the oscillatory breakdown between (i) the interfacing of normal linguistic cognition with the dynomic operations responsible for perception, and (ii) the interfacing of normal linguistic cognition with the dynomic operations responsible for conceptual understanding (Figure 2)

**Figure 2.**
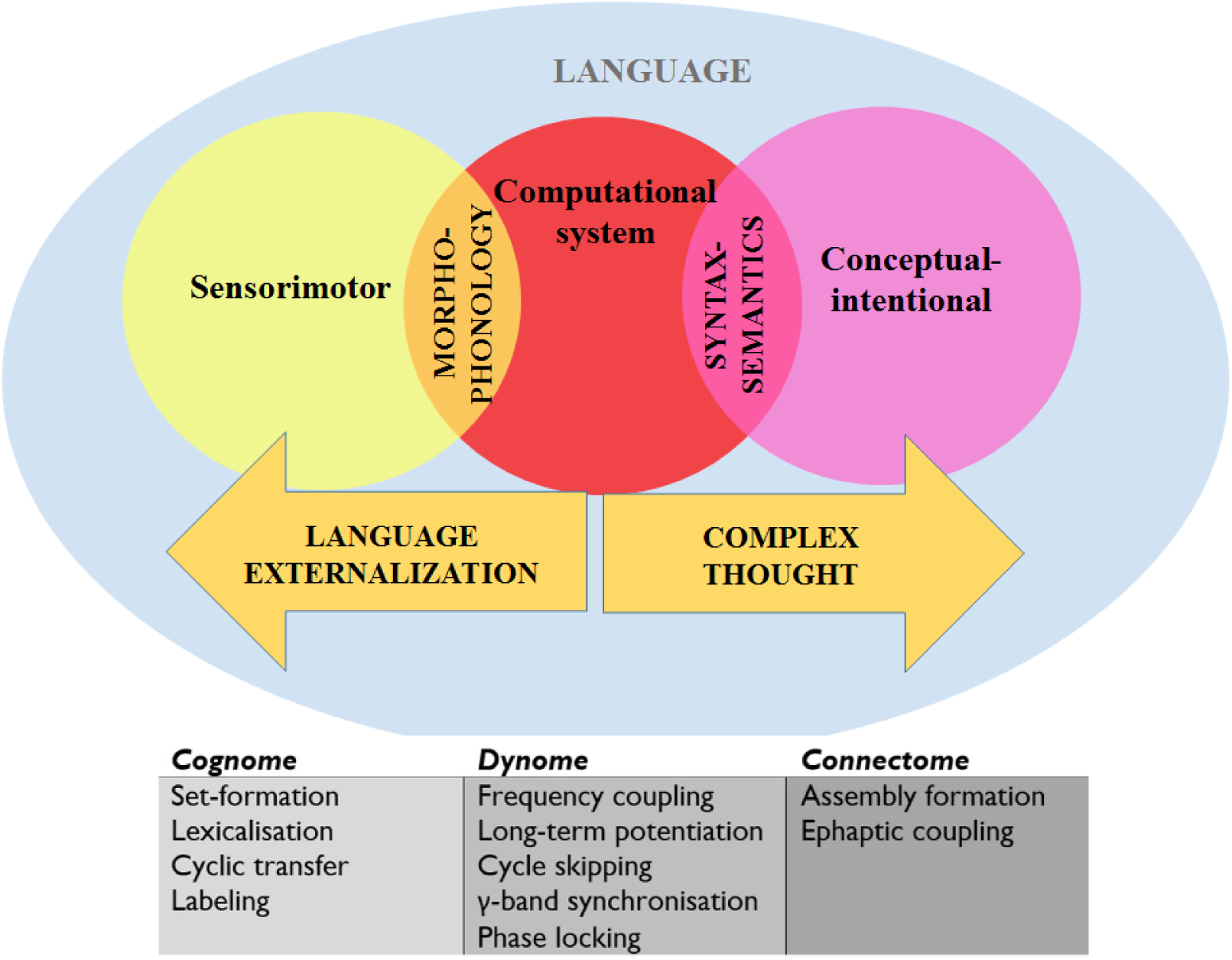
A schematic view of language representing the systems and interfaces of interest (above) and of the central operations implicated in the model of the cognome/dynome/connectome discussed in the paper. ‘Cognome’ refers to the operations available to the human nervous system (Poeppel, 2012); ‘dynome’ refers to brain dynamics (Kopell et al., 2014), and ‘connectome’ refers to the structural and functional interconnection patterns of the human brain (Rockland 2015).

Upon the emergence of human language’s uniquely recursive neural mechanisms, there also emerges a risk that it will over-generate and construct delusional propositional knowledge (‘I am Jesus’, etc.) which disturbs the balance between personal and non-personal understanding. Many of these problems may amount to disturbances in deictic shifts, a wholly grammatical phenomenon. These types of delusions generate novel thoughts detached from world knowledge – a human-specific ability which can be causally linked to modern, complex language. What exactly the neural mechanisms are which gave rise to these properties, and how they can be used to construct successful endophenotypes of schizophrenia, are the topics addressed in the next section.

## 3. From brain rhythmicity to language deficits in schizophrenia

Although schizophrenia was for a time deemed ‘the graveyard of neuropathology’ (Plum 1972) due to its unusually subtle neurophysiological markers, we believe that research in neuronal dynamics (particularly over the past half-decade) has the potential to carve a clearer image of the abnormally-developing brain. Neural oscillations in particular have the potential of allowing researchers to move beyond the still commonly discussed dopamine theories which point to the supersensitive dopamine receptors in the brains of schizophrenics (Owen et al. 1978) or the outdated ‘enlarged ventricles’ theory (Gattaz et al. 1991). Through oscillations, the brain becomes a self-organizing system whose capacities arise out of high-dimensional and usually nonlinear dynamics (Uhlhaas and Singer 2015). Oscillations play a central role in selectively enhancing neural assembly interconnectivity and information processing through the provision of spatio-temporal windows of enhanced or reduced patterns of excitability (Jensen et al. 2014, Weisz et al. 2014), and are consequently strong candidates for the origin of certain cognitive faculties.

Classic work such as Frith’s (1992: xi) monograph pointed to the ‘information processing abnormalities’ in schizophrenia, but a comprehensive account of their neurobiological causes has eluded researchers until the recent, burgeoning studies of neural oscillations. If the translational approach taken in Murphy (2015a) towards the brain dynamics of language is even approximately accurate, and if Hinzen and Rosselló (2015) are correct in claiming that linguistic disorganisation in schizophrenia “plays a more central role in the pathogenesis of this disease than commonly supposed”, then it is appropriate to inform our understanding of schizophrenia by focusing on the central role of brain rhythms in linguistic computation. If schizophrenia represents a breakdown in normal linguistic cognition, then we would expect to see disruptions in the model of brain dynamics of language processing outlined in Murphy (2015a) when examining the recent, burgeoning literature concerning the oscillatory profile of schizophrenics. This task will be the main focus of the current section.

To briefly summarise previous work, it was claimed in Murphy (2015a) that set-formation amounts to the *α* rhythm embedding cross-cortical *γ* rhythms, with *α* reflecting long-range cortical interactions (Nunez et al. 2001) and thalamo-cortical loop activity (Nunez and Srinivasan 2006). The syntactic operation of ‘Transfer’ (which ‘chunks’ constructed objects into short-term memory) was claimed to amount to the embedding of these *γ* rhythms inside the *θ* band, generated in the hippocampus. It was also claimed that labeling (maintaining an item in memory before coupling it with another, yielding an independent syntactic identity) amounts to the slowing down of *γ* to *β* before *β-α* coupling, likely involving a basal ganglia-thalamic-cortical loop. We will adopt these assumptions here when interpreting the rhythmic literature on schizophrenia (Figure 2).

Since schizophrenia, like other cognitive impairments, appears not to be the result of a locally delimited neural deficit but rather emerges from distributed impairments, neural oscillations and their role in flexible brain connectivity have recently become the target of research. Investigating the frequency and brain location of the neural oscillations involved in lexical processing in schizophrenia, Xu et al. (2013) conducted an MEG study in which patients discriminated correct from incorrect visually presented stimuli. This lexical decision task revealed that the patients, relative to healthy controls, showed abnormal oscillatory activity during periods of lexical encoding and post-encoding, particularly in the occipital and left frontal-temporal areas (see also Sun et al. 2014). Since a broad range of rhythms were implicated, we will avoid speculation about the specific operations impaired and instead suggest that the results imply familiar problems with semantic memory. However, the results did reveal reduced temporal lobe *α* and left frontal lobe *β* activity during lexical processing, suggesting difficulties in assigning lexical classes (labels) to items and successful categorization (findings corroborating the cartographic profile presented above, which included reduced activation during complex sentence processing at left superior frontal cortex). These findings are reminiscent of McClain’s (1983) claim that schizophrenic patients fail to spontaneously use semantic categories to memorize items during recall tasks (findings which directly relate to the pedantic speech documented above). These results corroborate the more general findings of reduced *α* and *β* in schizophrenia by Moran and Hong (2011) and Uhlhaas et al. (2008). A level of thalamocortical dysrhythmia was also detected by Schulman et al.’s (2011) MEG study; a discovery which bears on the claim that thalamocortical axons also likely play a role in language externalization (Boeckx and Benítez-Burraco 2014b). These suggested problems with the mechanisms responsible for phrase structure building also gain support from Ghorashi and Spencer’s (2015) findings that attentional load increases *β* phase-locking factor at frontal, parietal and occipital sites in healthy controls during a visual oddball task but not in schizophrenic patients (although this varied across individuals of different abilities), with the latter group having difficulty attending to and maintaining relevant objects in memory (perhaps as a result of their semantic memory deficits). *β*-generating circuits may well be responsible, then, for the types of computations attributed to them in Murphy (2015a).

An earlier MEG sentence presentation task by Xu et al. (2012) also found reduced *α* and *β* in left temporal-parietal sites, along with reduced *δ* at left parietal-occipital and right temporal sites and reduced *θ* at occipital and right frontal lobe sites, suggesting problems with phrase structure chunking; that is, problems with word movement and phrasal embedding, as attested above (see Ferrarelli et al. 2012). Schizophrenic patients also displayed reduced *δ* synchrony at left frontal lobe sites after sentence presentation, suggesting semantic processing dysfunctions. These findings are consistent with Hirayasu et al.’s (1998) MRI study of schizophrenic and bipolar individuals, which reported relatively reduced gray matter volumes in the left superior temporal gyrus for schizophrenics. Their results also give some support to the present hypothesis about chunking difficulties in schizophrenia, since they also reported reduced hippocampal volumes. Altogether, these studies are in agreement the findings of Hoffman et al. (1999), who suggested that the core schizophrenic deficit is not centred on attentional-perceptual cognitive processes, but rather verbal working memory (and, hence, difficulties with syntactic computation, given the ‘chunking’ nature of linguistic phrase structure building; see Narita 2014), mediated by oscillations generated in the hippocampus and left temporal regions (Murphy 2015a). Başar-Eroğlu et al. (2011) also documented reduced anterior *α* in response to simple auditory input, suggesting less efficient processing power.

Power and synchrony reductions in evoked *γ* have also been documented in chronic, first-episode and early-onset schizophrenia (Williams and Boksa 2010). Given the role of this band in feature binding and object representation (Uhlhaas et al. 2008) and its functional significance in the present dynomic model (Murphy 2015a), this suggests that schizophrenics have difficulties generating the correct category of semantic objects to employ in successful phrase structure building, as the behavioural results of lexical decision and related tasks appear to verify (likely explaining the features of delusions and formal thought disorder reviewed above). More recent studies appear to support this perspective. The amplitude of EEG *γ* was measured during phonological, semantic and visuo-perceptual tasks by Spironelli and Angrilli (2015). Schizophrenic patients, relative to normal controls, exhibited a significantly weaker hemispheric asymmetry across all tasks and reduced frontal *γ*. Ferrarelli et al. (2008) also found a decreased *γ* response in schizophrenic patients after TMS stimulation to the frontal cortex, suggesting an impaired ability to efficiently generate this rhythm. This is of particular significance given that *γ* amplitude has been shown to scale with the number of items held in working memory (Roux et al. 2012), and the limited phrase structure building and syntactic embedding capacities of schizophrenic patients would follow naturally from these results. Moreover, these cases of reduced *γ* may be the result of inhibitory interneuron malfunction (Lewis et al. 2012, Gonzalez-Burgos et al. 2015), and fMRI studies provide convergent evidence of deficits in the prefrontal cortex (e.g. Minzenberg et al. 2009) which can only be inferred via evoked potentials.

Recall also that the model of linguistic computation adopted here invokes a number of cross-frequency coupling operations. It is of interest, then, that schizophrenic patients showed higher γ-α cross-frequency coupling in Popov and Popova’s (2015) study of general cognitive performance, despite this co-varying with poorer attention and working memory capacities. The reason for this may be that the increased phase-amplitude-locking likely results in smaller ‘gamma pockets’ of working memory items (as Korotkova et al. 2010 argue on independent grounds) and hence low total *γ* power. In this instance, the size and order of working memory sequences outputted by the conceptual systems is not optimally compatible with the oscillopathic profile, leading to greater rhythmic excitability and yet inhibited linguistic functionality. Global rhythmicity is consequently disrupted due to unusually strong frontoparietal interconnectivity. We believe that this represents a genuine neural mechanism of an ‘interface’ between syntactically generated conceptual representations and external (memory) systems; a highly significant finding if corroborated by further experimental studies. Importing standard assumptions from generative grammar, we can think of the computational system as imposing its own conditions on the interfaces (Chomsky 1995). The shift in perspective to dynomic terms adopted here allows us to reformulate this such that the neural ensembles responsible for storing representations (lexical roots) used to construct phrases require particular phase-amplitude-locking levels in order for the interconnected regions coupled with them to ‘read off’ their content (figure 2).

Corroborating Angrilli et al.’s (2009) above hypothesis about schizophrenia being a failure of left-hemispheric phonological dominance, an MEG study of the oscillatory differences between bipolar disorder and schizophrenia revealed that schizophrenic patients showed delayed phase-locking in response to speech sounds in the left hemisphere, relative to bipolar individuals and normal controls (Oribe et al. 2010). This lack of left-hemispheric dominance may trigger confusion about internal and external voices and bring about a number of delusions, with language’s normal computational functioning being derailed. The left hypofrontality documented by Spironelli et al. (2011), with schizophrenic patients showing greater *δ* amplitudes over language-relevant sites (that is, greater functional inhibition), similarly point to a general functional deficit at the core memory sites of linguistic representations. It is also significant that the role attributed to *θ* in the present dynomic model gains support from the finding that this rhythm has greater amplitude in left superior temporal cortex during auditory hallucinations in schizophrenia (Ishii et al. 2000), as opposed to steady *θ* during resting state, with patients being seemingly incapable of regulating chunking operations. This perspective is complemented by an EEG study by Henshall et al. (2013) which revealed reduced interhemispheric coherence at auditory cortical areas during verbal hallucinations. *N*-methyl-D-aspartate (NMDA) hypofunction has been argued to modify the response of auditory cortex in schizophrenic patients to salient stimuli by attenuating the layer 4 *γ* band (Ainsworth et al. 2011). Given the identification of such dysrhythmias in schizophrenia, repetitive TMS (rTMS) could be used as a therapeutic intervention to modulate the oscillations responsible for the abnormal linguistic profile documented above, as has been done to improve performance on visual tasks (Farzan et al. 2012, Barr et al. 2013). The oscillopathic profile constructed here is presented in Table 1.

**Table 1.**
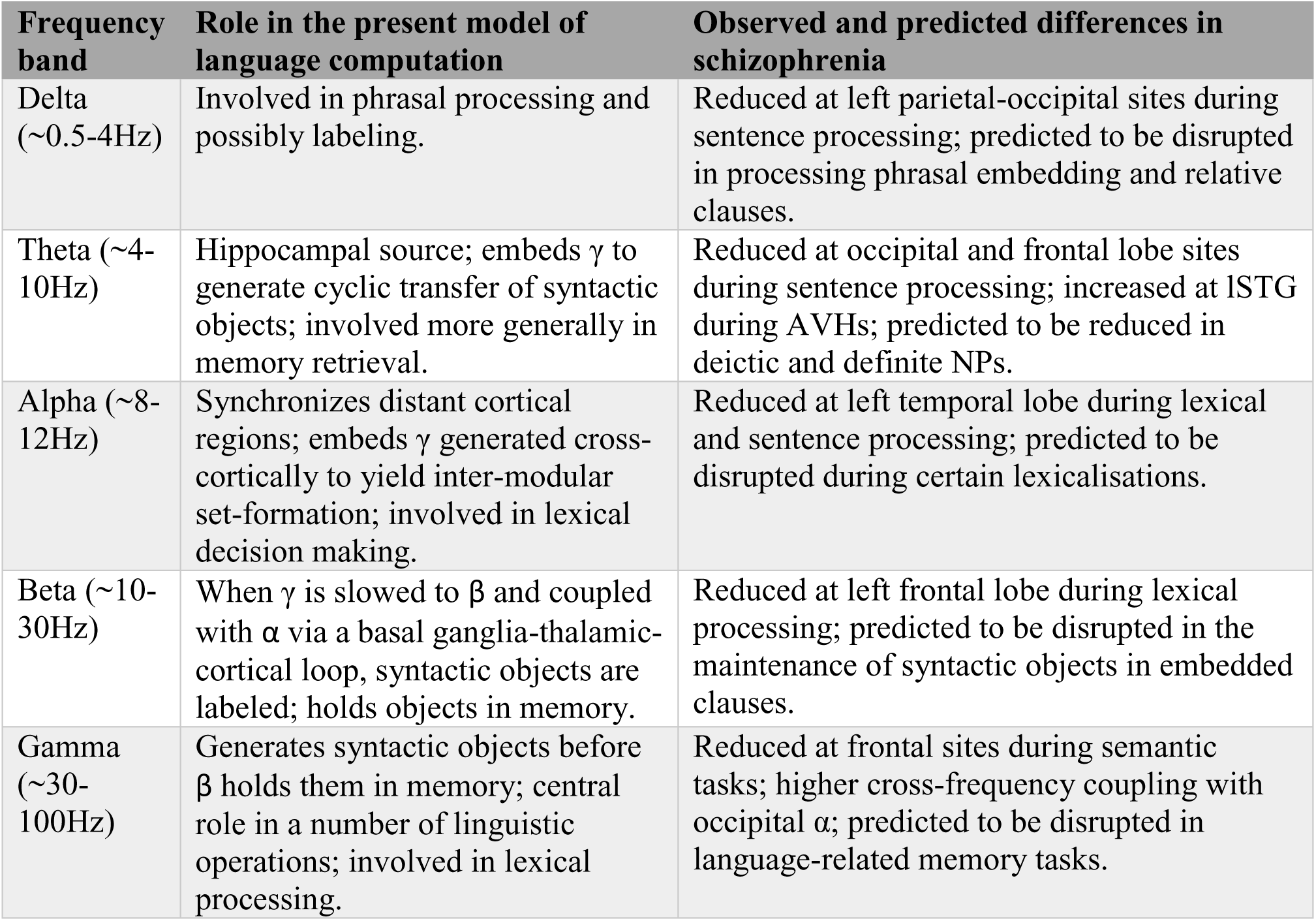
Summary of the present cognome-dynome model of linguistic computation and the observed differences in schizophrenia; lSTG denotes left superior temporal gyrus, AVH denotes auditory verbal hallucination.

We have so far posited a number of linking hypotheses between the cognome, dynome and connectome of schizophrenics. Our attention now turns to the genome and broader evolutionary considerations.

## 4. Schizophrenia-related genes and language evolution

As noted in the introduction, the set of candidate genes for schizophrenia has been growing over recent years. Several studies suggest that many of these genes map onto specific pathways and brain processes that are associated with susceptibility to the condition. Likewise, several candidates for language impairment in schizophrenia have been identified to date. Among them one finds *CNTNAP2* (Poot 2015), *SHANK3* (Guilmatre et al. 2014), *LYRM4* (Bozza et al. 2012), *TCF4* (Hui et al. 2015), and some of the genes located within the 3Mb region deleted in 22q11.2 deletion syndrome (Ousley et al. 2007); particularly, *COMT* (Egan et al. 2001; Malhotra et al. 2002), *PRODH* (Li et al. 2008) and *ZDHHC8* (Mukai et al. 2004). Interestingly, most of the genes related to cognitive dysfunction (and plausibly to language impairment) in schizophrenia map onto specific brain functions too. Hence, most of them are related to the homeostasis of neurotransmitters glutamate, dopamine, acetylcholine, and GABA (like *COMT*), but some of them encode ion transporters across the membrane (like *CAMK2A*, *CAMKK2*, or *CAMK4*) or are involved in corticogenesis (*NRG1, DISC1, RELN*) (see Papaleo et al. 2012 for details).

In the first part of this section, we will focus on genes specifically involved in the maintenance of the adequate balance between neuronal excitation and inhibition, and more generally, of brain rhythmicity, that are also candidate for schizophrenia. We will ask if the functions they perform helps understand the oscillopathic nature of the schizophrenic brain as outlined in the previous section, and, particularly, the language deficits that are characteristic of this condition. In the second part, we will review candidate genes for schizophrenia that may have played a relevant role in the evolution of modern cognition/language, with a special focus on brain connectivity and function.

As noted earlier in the introduction, brain oscillation patterns are highly heritable traits. Accordingly, we should expect that deviant the cognition of schizophrenics boils down to the oscillopathic activity of their brains resulting in part from the pathogenic variants of genes involved in brain function and rhythmicity. Among the most promising candidates one finds *ZNF804A*. This gene encodes a zinc finger binding protein important for cortical functioning and neural connectivity and involved in growth cone function and neurite elongation (Hinna et al. 2015). Schizophrenia risk polymorphisms of *ZNF804A* have been related to differences in performance in the domain of phonology, such as in reading and spelling tasks (Becker et al. 2012), but also in the domain of semantics, specifically in task evaluating category fluency (Nicodemus et al. 2014). Overall, these differences point to variances in memory processing, particularly in the visual domain (Hashimoto et al. 2010; Linden et al. 2013). On top of this, this gene has also been associated with verbal deficits in people with autism (Anitha et al. 2014). ZNF804A modulates hippocampal *γ* oscillations and, ultimately, the co-ordination of distributed networks belonging to the hippocampus and the prefrontal cortex (Cousijn et al. 2015), which are aspects known to be impaired in schizophrenia, as noted above (Uhlhaas et al. 2008; Godsil et al. 2013). Likewise, both NRG1 and its receptor ERBB4, which are strong candidates for schizophrenia (Hatzimanolis et al. 2013; Hou et al. 2014; Tost et al. 2014), enhance synchronized oscillations of neurons in the prefrontal cortex, known to be reduced in schizophrenia, via inhibitory synapses (Fisahn et al. 2009, Hou et al. 2014). Specifically NRG1 increases the synchrony of pyramidal neurons via presynaptic interneurons and the synchrony between pairs of interneurons through their mutually-inhibitory synapses (Hou et al. 2014). Risk polymorphisms of *NRG1* are associated with increased IQs as well as memory and learning performance, along with language in subjects with bipolar disorder (Rolstad et al. 2015). Moreover, risk alleles for the gene correlate with reduced left superior temporal gyrus volumes (a robust imaging finding in schizophrenia) (Tosato et al. 2012), a region related to language abilities (Aeby et al. 2013). Another gene of interest is *PDGFR*, which encodes the *β* subunit of the platelet-derived growth factor (PDGF) receptor, known to be involved in the development of the central nervous system. *Pdgfr-β* knocked-out mice show reduced auditory phase-locked *γ* oscillations, which correlates with anatomical (e.g. reduced density of GABAergic neurons in the amygdala, hippocampus, and medial prefrontal cortex), physiological (alterations of prepulse inhibition) and behavioral (reduced social behaviour, impaired spatial memory and problems with conditioning) hallmarks of schizophrenia (Nguyen et al. 2011, Nakamura et al. 2015). Interestingly, PDGFRA has been found to act downstream of FOXP2, the renowned ‘language gene’, to promote neuronal differentiation (Chiu et al. 2014) (more on *FOXP2* below).

Not surprisingly among the candidates for schizophrenia known to alter normal brain oscillation patterns, one finds several genes that encode ion channels. Genome-wide analyses (GWAs) have identified *CACNA1I* as one of the genes affecting sleep spindles in schizophrenics, a type of brain rhythm that recurs during non-rapid eye movement sleep and that constrains aspects of the thalamocortical crosstalk, impacting on sensory transmission, cortical plasticity, memory consolidation, and learning (Manoach et al. 2015). *CACNA1I* encodes a calcium channel and is abundantly expressed in the spindle generator of the thalamus. Likewise *CACNA1C* encodes the alpha 1C (*α*1C) subunit of the Cav1.2 voltage-dependent L-type calcium channel, a calcium channel involved in the generation of *β* to *γ* waves during wakefulness and rapid eye movement (REM) sleep, and ultimately in sleep modulation; all of which are aspects known to be altered in schizophrenics (Kumar et al. 2015). Intriguingly, *CACNA1C* is related to semantic (but not lexical) verbal fluency in healthy individuals; conversely, risk alleles of this gene correlate with lower performance scores, seemingly accounting for the non-fluent verbal performance of schizophrenics (Krug et al. 2010).

Proteins associated with ion channels are also worth consideration. *DPP10* is a gene associated with schizophrenia and bipolar disorder (Djurovic et al. 2010) which encodes a membrane protein that binds specific potassium channels and modifies their expression and biophysical properties. Similarly, CNTNAP2 is a protein associated with K% voltage-gated channels in the axon initial segment of pyramidal cells in the temporal cortex, that are mostly innervated by GABAergic interneurons (Inda et al. 2006). *CNTNAP2* is a candidate for several types of language disorders, including child apraxia of speech (Worthey et al. 2013), dyslexia (Peter et al., 2011), SLI (Newbury et al. 2011), language delay and language impairment (Petrin et al. 2010, Sehested et al. 2010). CNTNAP2 additionally affects language development in the normal population (Whitehouse et al. 2011, Whalley et al. 2011, Kos et al. 2011), apparently because of its effects on brain connectivity and cerebral morphology (Scott-Van Zeeland et al. 2010, Tan et al. 2010, Dennis et al. 2011) and dendritic arborization and spine development (Anderson et al. 2012). *CNTNAP2* is also a target of FOXP2 (Vernes et al. 2008).

Not surprisingly, genes encoding neurotransmitter receptors have been recurrently related to abnormal brain oscillation patterns in schizophrenia. *HTR1A* encodes the receptor 1A of serotonin and modulates hippocampal *γ* oscillations, seemingly impacting on behavioural and cognitive functions, such as learning and memory linked to serotonin function (Johnston et al. 2014). Similarly, receptors of NMDA, particularly those containing the subunit NR2A, encoded by *GRIN2A*, are known to be reduced in fast-firing interneurons in schizophrenics, which plays a critical role in *γ* oscillation formation; a blockade of NR2A-containing receptors gives rise to strong increases in *γ* power and a reduction in low-frequency *γ* modulation (Kocsis, 2012). More generally, mutations in *GRIN2A* cause epilepsy-aphasia spectrum disorders, including Landau-Kleffner syndrome and continuous spike and waves during slow-wave sleep syndrome (CSWSS), in which speech impairment and language regression are prominent symptoms (Carvill et al. 2013, Lesca et al. 2013). The gene has been related as well to rolandic epilepsies, the most frequent epilepsies in childhood, in which cognitive, speech, language and reading problems are commonly observed (Dimassi et al. 2014). Speech problems linked to *GRIN2A* mutations include imprecise articulation, impaired pitch and prosody, and hypernasality, as well as poor performance on maximum vowel duration and repetition of monosyllables and trisyllables, resulting in lifelong dysarthria and dyspraxia (Turner et al. 2015). Finally, cannabinoid-1receptor, encoded by *CNR1*, modulates *θ* and *γ* oscillations in several areas of the brain, including the hippocampus, impacting on sensory gating function in the limbic circuitry (Hajós et al. 2008). The gene has also been linked to cases of complete absence of expressive speech (Poot et al. 2009).

Finally, we wish to highlight one of the strongest candidates for schizophrenia, namely, *DISC1* which encodes a protein involved in neurite outgrowth, cortical development and callosal formation (Brandon and Sawa 2011; Osbun et al. 2011). In hippocampal area CA1 of a transgenic mouse that expresses a truncated version of *Disc1* mimicking the schizophrenic genotype, *θ* burst-induced long-term potentiation (and ultimately, long-term synaptic plasticity) is altered (Booth et al. 2014). The ability of DISC1 to regulate excitatory-inhibitory synapse formation by cortical interneurons depends on its inhibitory effect on NRG1-induced ERBB4 activation and signalling (more on which below), ultimately effecting the spiking interneuron-pyramidal neuron circuit (Seshadri et al. 2015). *DISC1* is also a target of FOXP2 (Walker et al. 2012)

A more systematic account of genes relating to language deficits (and aberrant patterns of brain oscillations linked to language deficits) in schizophrenia emerges from evolutionary studies aimed to explore the genetic basis of our species-specific ability to learn and use language, i.e. our language-readiness. Many authors have pointed to the stable prevalence of schizophrenia across cultures and epochs, and have explored the reasons why susceptibility genes for this condition, otherwise tenuous or absent in great apes, have been preserved in the human genepool. It seems that what is disadvantageous at the individual level may be neutral or even yield some advantage at the group level in specific social contexts (see Brüne 2004 and Pearlson and Folley 2008 for discussion). More specifically, it has been hypothesised that schizophrenia candidate genes were involved in the evolution of the human brain and that the processes they contributed to improving are identical to those impaired in schizophrenics. For example, the human prefrontal cortex, which is responsible for many human-specific cognitive abilities, is differently organized in humans compared to great apes as a result of a recent reorganization of the frontal cortical circuitry; at the same time, these circuits are impaired in schizophrenia and other psychiatric and neurological conditions (Teffer and Semendeferi 2012). Similarly, reduced levels of dopamine are observed in schizophrenics, but we know that this neurotransmitter is important for social behaviour and is involved in transmission between areas that play key roles in cognitive and affective functions (see Yamaguchi et al. 2015 for review). Regarding the connection between language evolution and schizophrenia, Arbib and Mundhenk (2005) have argued that the mirror system hypothesis may account for both the evolution of the language-ready brain and the schizophrenic phenotype. The mirror system hypothesis claims that primate mirror neurons, which fire both when the animal manipulates an object and when it sees another conspecific manipulating it, provided the scaffolding for imitation abilities involved in language acquisition. Interestingly, schizophrenics show a spared ability to generate actions, whether manual or verbal, but they lack the ability to attribute the generation of that action to themselves. More specifically, it has been suggested that schizophrenia is the ‘price we paid for language’ (Crow 1997). Accordingly, Crow and colleagues have hypothesised that schizophrenia represents an extreme of variation of hemispheric specialisation and that a single genetic mechanism involving both the X and Y chromosomes, that was modified during recent human history, can account for this variation because it generates epigenetic diversity related to both the species capacity for language and the predisposition to psychosis (Crow 2008).

Nonetheless, when it comes to testing the hypothesis that the same genes that cause schizophrenia (and other cognitive diseases) refined the brain processes that led to modern cognition and language, contradictory results have been obtained. Concerning the protein-coding regions of genes associated to psychiatric disorders Ogawa and Wallender (2014) did not find evidence of differential selection in humans compared to non-human primates, although elevated dN/dS was observed in primates and other large-brained taxa like cetaceans. However, recent analyses based on large GWAs of schizophrenia and data of selective sweeps in the human genome compared to Neanderthals suggest that brain-related genes showing signals of recent positive selection in anatomically-modern humans (AMHs) are also significantly associated with schizophrenia (Srinivasan et al. 2015), supporting the view that schizophrenia is a by-product of the changes in the human brain that led to modern cognition and language. Interestingly, among the loci highlighted by Srinivasan et al. (2015), one finds genes related to language development, language impairment, and language evolution (see also Figure 4). Among them, we wish highlight: *FOXP1*, *GATAD2B*, *MEF2C*, *NRG3*, *NRXN1*, and *ZNF804A. FOXP1* encodes an interactor of FOXP2 (Li et al. 2004). *FOXP1* is expressed in areas relevant to cortico-laryngeal connections (Inoue et al. 2008) and its mutations cause language impairment, intellectual disability, and autism (Hamdan et al. 2010; Sollis et al. 2015). *FOXP1* is mentioned among the top five percent regions showing signals of positive selection in AMHs (Green et al. 2010). *GATAD2B* encodes a zinc protein involved in chromatin modification and regulation of gene expression; mutations in *GATAD2B* impact synaptic growth and function (Willemsen et al. 2013) and have been related to mental retardation, intellectual disability, and learning problems (De Ligt et al. 2012; Hamdan et al. 2014; Roberts et al. 2014), and specifically, to limited speech (Willemsen et al. 2013). *MEF2C* encodes a trans-activating and DNA binding protein involved in early neurogenesis, neuronal migration, and differentiation. Mutations in *MEF2C* cause absent speech, severe mental retardation, and epilepsy (Bienvenu et al. 2013). *MEF2C* is a target of FOXP2 in the basal ganglia (Spiteri et al. 2007). *NRG3* is a promising candidate for atypical neurodevelopmental outcomes (including cognitive anomalies and abnormal infant behaviour) that may affect preterm infants in absence of rare genetic diseases (Blair et al. 2015). Deletions and duplications involving *NRG3* give rise to speech delay (van Bon et al. 2011). In conjunction with NRG1 and their receptor ERBB4 (reviewed below), NRG3 regulates the migration of GABAergic interneurons from ganglionic eminences to their cortical targets (Li et al. 2012). Finally, *NRXN1* encodes one of the largest known neurexins, a presynaptic cell adhesion molecule important for synaptic activity, neuritogenesis, and neuronal network assembly related to neocortical development (Südhof, 2008; Gjørlund et al. 2012; Jenkins et al. 2015). Mutations in *NRXN1* impact speech severely, although give rise to mild motor delay only (Zweier 2012).

**Figure 4.**
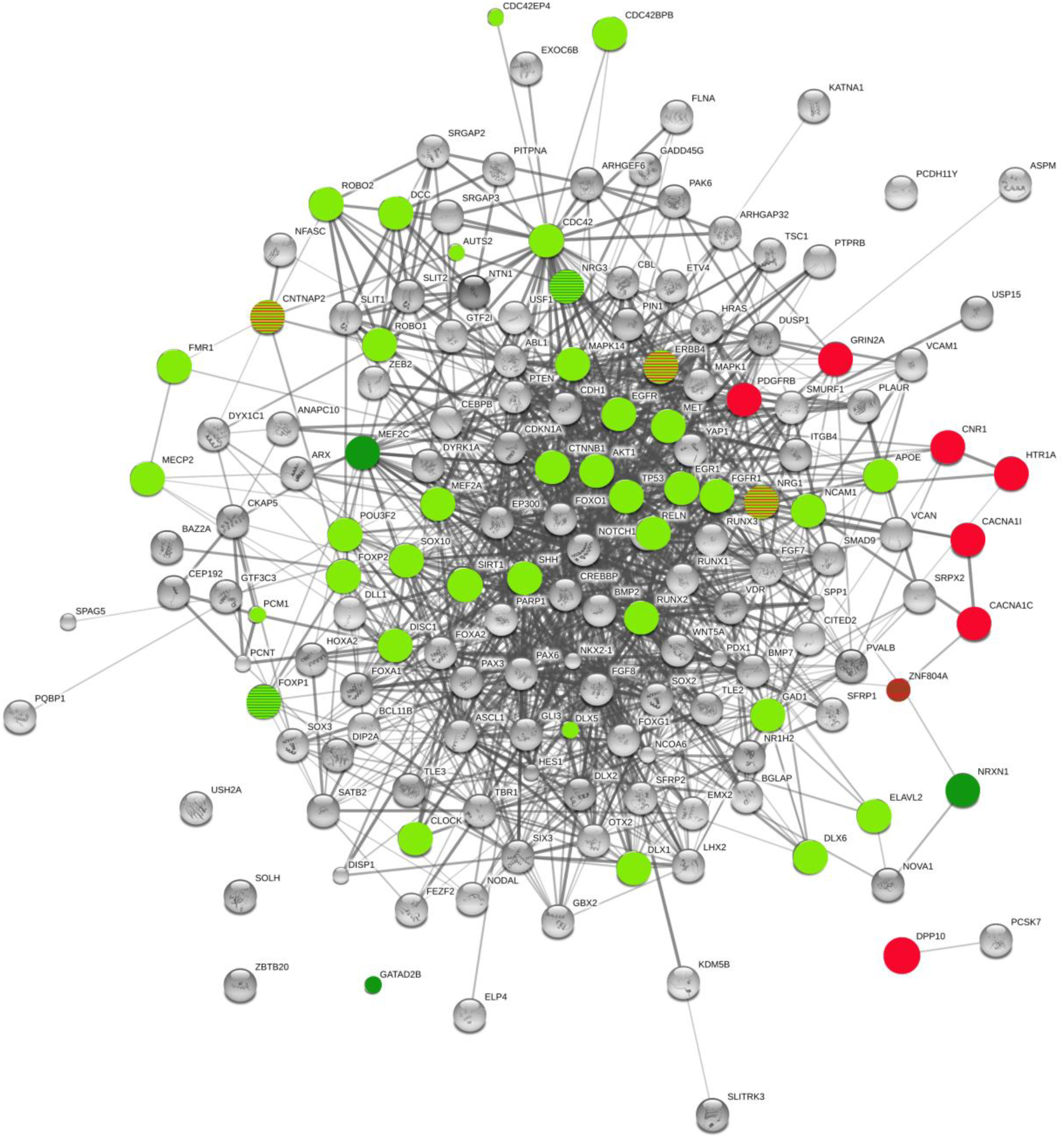
Functional links predicted by String 10 among candidates for the evolution of language, candidate genes for schizophrenia, and genes important for brain rhythmicity. Candidate genes for the evolution of language (as posited by Boeckx and Benítez-Burraco 2014a,b, and Benítez-Burraco and Boeckx 2015) that are also candidates for schizophrenia are colored in light green (otherwise they appear in grey). Genes related to brain rhythms are colored in red, but they appear stripped in red and light green if they belong to any of the interactomes important for language evolution. Candidate genes for schizophrenia showing signals of positive selection in AMHs according to Srinivasan et al. (2015) are colored in dark green, but they appear stripped in dark and light green if they also belong to the list of candidates for language evolution. Only one gene, namely, ZNF804A, is stripped in red and dark green, meaning that it is both related to brain oscillations and has been selected in AMHs, although we have not described it yet as part of the putative interactome for the language-ready brain. The graph displays four candidates for language-readiness and schizophrenia that have not been discussed in the main text: FGFR1, FMR1, MECP2, and TGF. Stronger associations between proteins are represented by thicker lines. The medium confidence value was .0400 (a 40% probability that a predicted link exists between two enzymes in the same metabolic map in the KEGG database: http://www.genome.jp/kegg/pathway.html). String 10 predicts associations between proteins that derive from a limited set of databases: genomic context, high-throughput experiments, conserved coexpression, and the knowledge previously gained from text mining (Szklarczyk et al. 2015). This is why the figure does not represent a fully connected graph (evidence for additional links are provided in the main text). Importantly, the diagram only represents the potential connectivity between the involved proteins, which has to be mapped onto particular biochemical networks, signaling pathways, cellular properties, aspects of neuronal function, or cell-types of interest that can be confidently related to aspects of language development and function (although see Table 2 and main text for some concerns regarding schizophrenia).

**Table 2.**
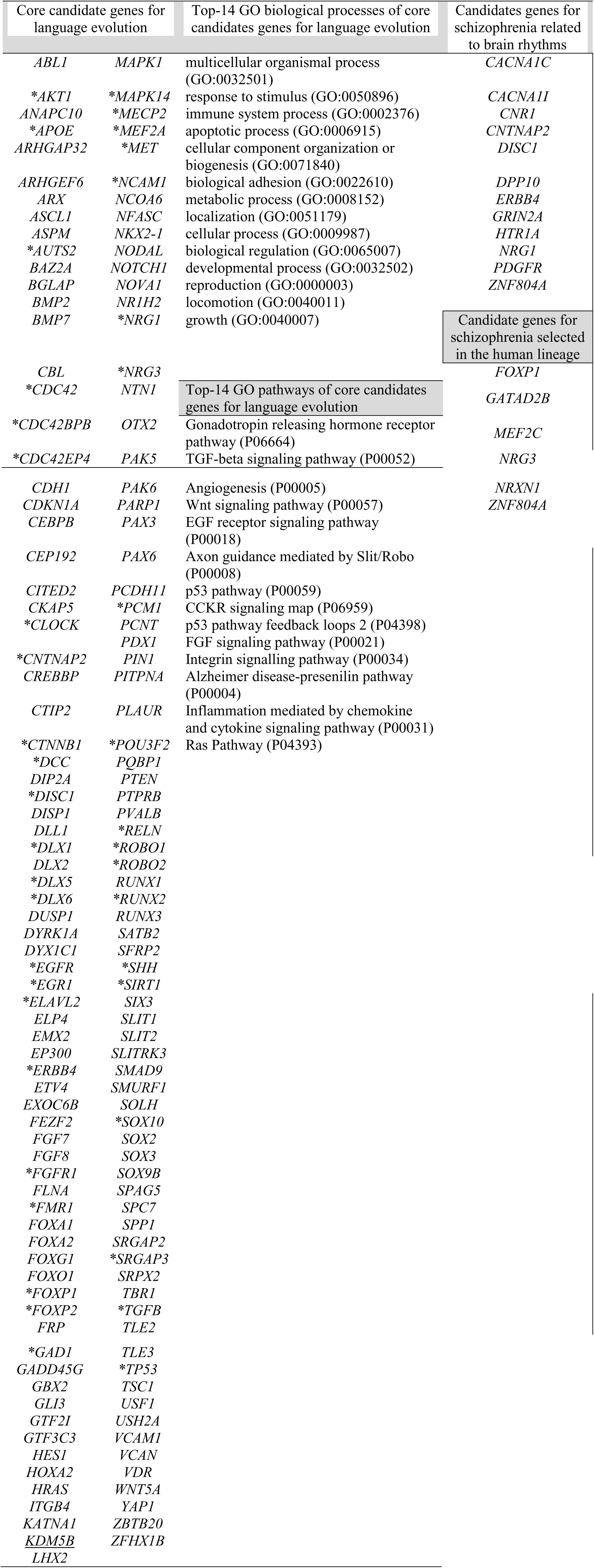
Genes discussed in Section 4. The first column contains core candidates for the evolution of language as posited by Boeckx and Benítez-Burraco (2014a,b) and Benítez-Burraco and Boeckx (2015). Candidates for schizophrenia are marked with an *. The second column provides a GO classification of these genes according to Panther (http://pantherdb.org; only the top-25 functions after a Bonferroni correction have been included. The last column comprises genes that have been related to brain rhythms (above) and candidate genes for schizophrenia showing signals of positive selection in AMHs according to Srinivasan et al. (2015) (below), highlighted here as potential new candidates for language evolution.

It should be highlighted that some of the genes involved in brain rhythmicity reviewed above also show differences in the human lineage. *DPP10* shows signals of differential expression in the human brain compared to primates. Hence, the gene displays differences in the methylation patterns of cis-regulatory regions affecting transcription start sites, as well as human-specific higher order chromatin structures indicative of human-specific gene expression patterns and networks; additionally, sequences at DPP10 show regulatory motifs absent in archaic hominins and signals of strong selection in modern human populations (Shulha et al. 2012). Likewise, DISC1 interacts with PCNT, mentioned by Green et al. (2010) as being amongst the proteins that show non-synonymous and non-fixed changes compared to Neanderthals. PCNT is a protein of the centrosome that has been related to dyslexia (Poelmans et al. 2009). Finally, the human CNTNAP2 protein bears a fixed change (I345V) compared to the Denisovan variant (Meyer et al. 2012) and it is related in addition to NFASC, a protein involved in postsynaptic development and neurite outgrowth (Kriebel et al. 2012) which also shows a fixed change (T987A) in AMHs compared to Neanderthals/Denisovans (Pääbo, 2014, Table S1).

A more comprehensive view of the connection between schizophrenia and language (evolution) is provided by recent analysis of the changes that prompted the emergence of our language-readiness. As pointed out in the introduction, this ability is rooted in a recently-evolved ability to form cross-modular concepts (known to be affected in schizophrenia) and it seemingly resulted from the changes in brain wiring linked to the globularization of the AMH skull that habilitated a new neuronal workspace (see Boeckx and Benítez-Burraco 2014a for details, and also Murphy 2015c). In a series of related papers, we have put forth a list of tentative candidates for globularization and language-readiness (Boeckx and Benítez-Burraco 2014a,b; Benítez-Burraco and Boeckx 2015; see Table 2 and Figure 4). The list encompasses genes involved in bone development, brain development (specifically of GABAergic neurons), and more generally, brain-skull cross-talk, like *RUNX2*, some DLX genes (including *DLX1*, *DLX2*, *DLX5*, and *DLX6*), and some *BMP* genes (like *BMP2* and *BMP7*). It also includes genes that regulate subcortical-cortical axon pathfinding and that are involved in the externalization of language (such as *FOXP2*, *ROBO1*, and the genes encoding the SLITs factors). Finally, it also comprises genes connecting the former two interactomes, including *AUTS2* and some of its partners (Figure 3). Many of these genes show differences with extinct hominin species, particularly, with Neanderthals/Denisovans, which affect their regulatory regions, their coding regions, and/or their methylation patterns (see Boeckx and Benítez-Burraco 2014a,b; Benítez-Burraco and Boeckx 2015 for details). Interestingly, among the genes highlighted by Benítez-Burraco and Boeckx we have found many candidates for schizophrenia (Table 2 and Figure 4). In the last part of this section we will briefly discuss the most relevant of these genes.

**Figure 3.**
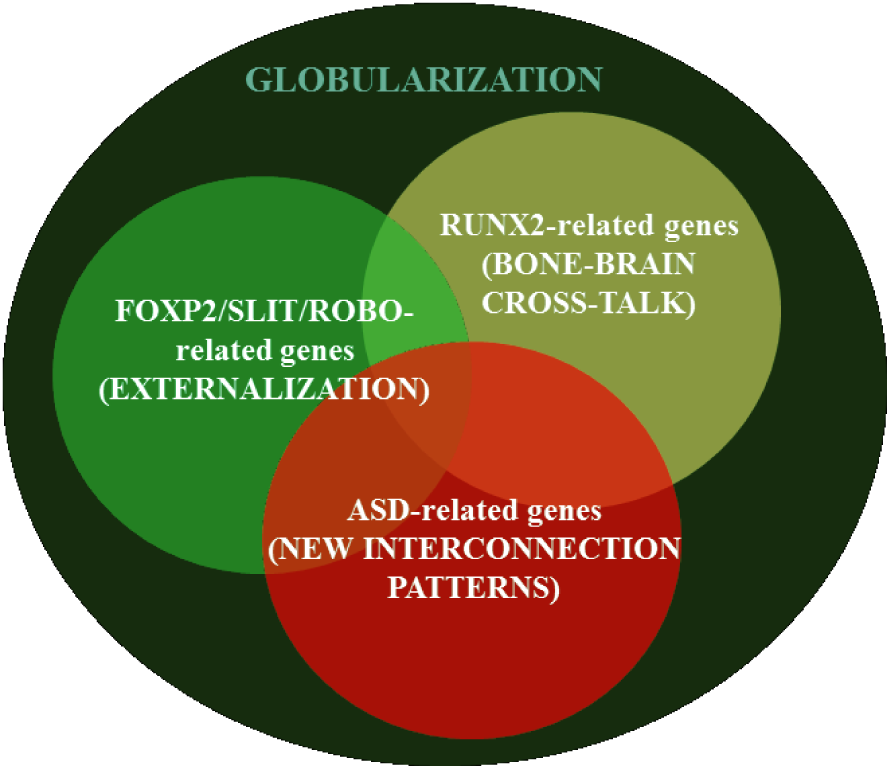
Three putative gene networks that may account for the emergence of language-readiness in our species. As noted in the main text, all of them include candidate genes for schizophrenia (based on Boeckx and Benítez-Burraco, 2014a,b and Benítez-Burraco and Boeckx, 2015a).

Regarding the genes clustered around RUNX2, we wish to note that *RUNX2* is listed among the genes associated with GAD1-dependent GABAergic dysfunction in schizophrenia (Benes et al. 2007). GAD1 regulatory network is important for the normal development of GABAergic neurons within the hippocampus (Pleasure et al. 2000, Ruzicka et al. 2015) and *GAD1* itself is a strong candidate for schizophrenia (Mitchell et al. 2015). Other genes important for globularity and the emergence of our language-ready brain besides *RUNX2* interact with GAD1, including *FOXP2*, *DLX1*, and *DLX2*. Importantly, the promoter region of *RUNX2* shows strong signals of a selective sweep in AMHs (Green et al. 2010). Moreover, the interaction between RUNX2 and VDR (the 1*α*,25-dihydroxyvitamin D3 receptor) regulates the expression of both *SPAG5* and *SRGAP3* (Stephens and Morrison 2014). *SPAG5* has been selected in AMHs (Green et al. 2010) and encodes an interactor of the isoform B of USH2A (Kersten et al. 2012), the main candidate for Usher syndrome, a condition involving combined deaf-blindness and occasional schizophrenia-like symptoms (Domanico et al. 2012; see Leivada and Boeckx 2014 for detailed discussion). In turn, *SRGAP3* is related to both schizophrenia (Wilson et al. 2011; Waltereit et al. 2012) and severe mental retardation and absence of speech (Endris et al. 2002). Interestingly, one interactor of SRGAP3 during neuronal differentiation and neurite outgrowth, namely SRGAP2 (Ma et al. 2013), has been duplicated three times in humans (Sudmant et al. 2010). Both SRGAP2 and SRGAP3 interact with ROBO1 and affect the SLIT/ROBO pathway (Wong et al. 2001), important for the externalization of language, as noted above (see Boeckx and Benítez-Burraco 2014b for details). RUNX2 also interacts with *APOE* (Kuhlwilm et al. 2013), a gene related to encephalization and cognitive development in our clade (Bufill and Carbonell 2006) and part of the Reelin signalling cascade related to cognitive dysfunction in schizophrenia, including verbal memory deficits (Verbrugghe et al. 2012; Li et al. 2015). *NCAM1*, which encodes a protein involved in axonal and dendritic growth, synaptic plasticity, and cognition, is a potential target of RUNX2 too (Kuhlwilm et al. 2013), but also of FOXP2 (Konopka et al. 2009). *NCAM1* has been related to schizophrenia (Vawter et al. 2001; Atz et al. 2007) and working memory performance (Bisaz et al. 2013). Interestingly, it interacts with VCAM1, a protein that shows a fixed change (D414G) in AMHs compared to Neanderthals/Denisovans (Pääbo, 2014, Table S1). VCAM1 is involved in cell adhesion in the subventricular zone (Kokovay et al. 2012). In turn, *VCAM1* is upregulated by CLOCK (Gao et al. 2014), a circadian gene associated to schizophrenia (Zhang et al. 2011; Jung et al. 2014), and an interactor of RUNX2 (Reale et al. 2013). VCAN is also functionally linked to EGFR, another of RUNX2’s targets (Kuhlwilm et al. 2013) and a candidate for schizophrenia too (Benzel et al. 2007), a link which reinforces the view that ERBB and NRG families are causative factors of the disease, as noted before. Among the genes belonging to the RUNX2 network we wish also highlight one finds *DLX1*, *DLX5*, and *DLX6*. Decreased expression of *DLX1* in the thalamus has been observed in schizophrenics (Kromkamp et al. 2003). Abnormal configuration of thalamic circuits is a hallmark of the disease, whereas changes in the thalamus are expected to have contributed to our mode of cognition (see Boeckx and Benítez-Burraco 2014a for details). DLX5 and DLX6 regulate GABAergic interneuron development (Cobos et al. 2006). Importantly, Dlx5/6(+/-) mice show and abnormal pattern of *γ* rhythms resulting from abnormalities in GABAergic interneurons, particularly fast-spiking interneurons, which impact on their cognitive flexibility (Cho et al. 2015). On the whole, the genes highlighted above are primarily related to the specification, migration and interconnection of GABAergic neurons within the forebrain, to skull morphogenesis and to thalamic development, all of them aspects known to be impaired in schizophrenia. This circumstance reinforces the view that globularization was brought about by changes in genes that are involved in schizophrenia when mutated.

Regarding the network centered around FOXP2 and the ROBO/SLIT factors, we wish to mention that *FOXP2* has been recurrently associated to schizophrenia (Li et al. 2013) and to some of the changes observed in the brain of schizophrenics, including a reduction of grey matter in areas involved in language processing that may contributed to the verbal hallucinations that are a hallmark of the disease (Španiel et al. 2011). As noted above several targets of FOXP2 are related to schizophrenia (*CNTNAP2*, *DISC1*, *MEF2C*). Also some of its effectors are related to the disease. For example, sequence and copy number variations affecting *POU3F2* have been found in subjects with schizophrenia (Huang et al. 2005; Potkin et al. 2009). Importantly, the AMH POU3F2 is less efficient than the Neanderthal version in activating transcription of *FOXP2* (Maricic et al. 2013). POU3F2 regulates dopamine and serotonin synthesis (Nasu et al. 2014) and neuronal migration and identity in the neocortex (McEvilly et al. 2002; Sugitani et al. 2002). Likewise, FOXP2 regulates *MET* (Mukamel et al. 2011), a gene that influences schizophrenia risk and neurocognition (Burdick et al. 2010). Interestingly, *FOXP2* and some other candidates for schizophrenia reviewed above, like *CNTNAP2* and *DLX1*, are enriched ELAVL2 target genes (Konopka et al. 2012). *ELAVL2* encodes a splicing factor involved in cortical neurogenesis whose expression pattern has changed in humans (Konopka et al., 2012), and it is a candidate for schizophrenia too (Yamada et al. 2011). Likewise both *ROBO1* and *ROBO2*, core components of our network, have been proposed as schizophrenia-candidate genes (Benes et al. 2009, Potkin et al. 2009, 2010). Both genes are involved in thalamocortical axon development, which represent the major input to the neocortex, and modulate cognitive functions, consciousness and alertness (López-Bendito et al. 2007; Marcos-Mondéjar et al. 2012). Both genes are differentially expressed in areas important for singing in adult male zebra finches (Wang 2011). In humans, *ROBO1* has been associated with dyslexia and speech sound disorder (Hannula-Jouppi et al. 2005; Mascheretti et al. 2014), whereas *ROBO2* has been associated with expressive vocabulary growth in the normal population (St Pourcain et al. 2014), and linked to dyslexia (Fisher et al. 2002) and speech-sound disorder and reading (Stein et al. 2004).

Other ROBO/SLIT-related genes that belong to our network and that are also candidates for schizophrenia are *ABL1*, *AKT1*, *CTNNB1*, *DCC*, *EGR1*, *MAPK14*, and *PCM1. ABL1* is involved in cell differentiation, division, and adhesion important for the regulation and/or the activation of auditory networks within the thalamus (Habib et al. 2013) and is differentially expressed in the hippocampus of schizophrenics (Benes et al. 2009). *AKT1* is involved in neuronal survival and bone formation (Dudek et al. 1997; Peng et al. 2003). In humans mutations in *AKT1* have been associated to schizophrenia (Emamian et al. 2004) and Proteus syndrome (Cohen 2014). *CTNNB1*, related to schizophrenia (like other components of the Wnt/β-catenin pathway) (Levchenko et al. 2015), interacts with *PCDH11X/Y*, the gene pair that has undergone accelerated evolution in our lineage (Williams et al. 2006) and that has been linked to language acquisition delay (Speevak and Farrell 2011) and to schizophrenia and language evolution, as noted above (see Crow 2013 for discussion). *DCC* is involved in thalamocortical axon projections and the organization of dopaminergic circuits within the cortex (Braisted et al. 2000; Grant et al. 2007). DCC contributes to the genetic basis behind individual differences in susceptibility to schizophrenia (Grant et al. 2007; Grant et al. 2012). Importantly, an hCONDEL (shared with Neanderthals) exist in a region upstream of *DCC* (McLean et al. 2011). *EGR1* is found differentially expressed in the prefrontal cortex of schizophrenics (Pérez-Santiago et al. 2012). This gene encodes a transcription factor involved in neuronal plasticity and memory consolidation (Veyrac et al. 2014). EGR1 downregulates *PLAUR* (Matsunoshita et al. 2011), a target of FOXP2 (Roll *et al*. 2010) which encodes an effector of SRPX2, another of FOXP2 targets (Royer-Zemmour et al. 2008) and a candidate for rolandic epilepsy and speech dyspraxia (Roll et al. 2006). *MAPK14* encodes an interactor of both ABL1 and AKT1 involved in cellular proliferation and differentiation, and it is also a candidate for brain changes in schizophrenia (Onwuameze et al. 2013). Finally, *PCM1*, which encodes a centrosome protein that interacts with SLIT1 and that is necessary for neuronal migration, shows a differential expression in mammalian vocal learners (Wang 2011). PCM1 also interacts with DISC1 in the centrosome, mimicking its effects on neural migration and cortical development (Kamiya et al. 2008). On the whole, these genes are prominent signatures of vocal learning, important for the externalization component of the language-ready brain, which is also impaired in schizophrenia, as described in sections 2 and 3.

Regarding the genes clustered around *AUTS2*, we wish to highlight that AUTS2 itself, though a strong candidate for autism, has been recently associated with the disease (Zhang et al. 2014). The first half of *AUTS2* displays the strongest signal of positive selection in AMHs compared to Neanderthals and contains several human accelerated regions which include enhancers that seem to be active in the brain (Green et al. 2010; Oksenberg et al. 2013). AUTS2 interacts with many proteins involved in brain development and function that are encoded by candidate genes for several neurodevelopmental disorders affecting cognition and language (reviewed by Oksenberg and Ahituv 2013), including RELN (mentioned above) and TBR1. TBR1 is a partner of DYRK1A, encoded by a gene that contains a region showing signals of strong selection in AMHs (Green et al. 2010) and whose mutations affect speech abilities (Van Bon et al. 2011; Courcet et al. 2012). DYRK1A regulates *GAD1* (Souchet et al. 2014) and it is important for the control of balance between excitation and inhibition in the brain and for synaptogenesis and synaptic plasticity (and, ultimately, for learning and memory) (Hämmerle et al. 2003; Souchet et al. 2014). Moreover, DYRK1A directly phosphorylates SIRT1 and also promotes deacetylation of TP53. Both *SIRT1* and *TP53* are candidates for schizophrenia (Ni et al. 2005; Kishi et al. 2011; Wang et al. 2015). Interestingly, SIRT1 is an effector of several genes under selection in modern populations that show non-fixed changes in their coding regions compared to Neanderthals and Denisovans, like *BAZ2A* and *NR1H2* (Prüfer et al. 2014). *SIRT1* is functionally related to *MEF2A* too (Gracia-Sancho et al., 2010), an important gene implicated in differences between human and chimpanzee prefrontal cortex development (Liu et al. 2012) and that shows signals of recent positive selection (Somel et al. 2013). According to Liu and colleagues (2012), these differences may account for the presumed faster cortical synaptic development in Neanderthals. Notably, a binding site for MEF2A has been linked to formal thought disorder (Thygesen et al. 2015). Likewise, TP53 exhibits a non-fixed change (P72R) compared to Neanderthals/Denisovans (Paskulin et al. 2012) and the expression pattern of the human gene differs from the patterns observed in other primates (Konopka et al. 2012). Risk alleles for TP53 seem to contribute to the reduced metabolic activity and the reduced white matter volumes observed in the frontal lobe of schizophrenics (Molina et al. 2011). On the whole, these changes seemingly contributed to the refinement of the changes that brought about modern cognition and enhanced speech abilities in humans.

Some other genes relevant for brain function that show changes that occurred after the split between AMHs and Neanderthals/Denisovans, and that reinforce the links between the three sets of genes highlighted above, are candidates for schizophrenia. We will focus only on those belonging to the CDC42 signaling pathway and the SHH-GLI signaling pathway. Firstly, *CDC42* is required for proper cortical interneuron migration (Katayama et al. 2013). Some risk polymorphisms for schizophrenia reduce the expression of *CDC42* (Gilks et al. 2012). Specifically, the downregulation of the gene in the dorsolateral prefrontal cortex appears to contribute to the reduction of dendritic spines on pyramidal cells and, ultimately, to the cognitive dysfunction characteristic of the disease (Datta et al. 2015). For our purposes, it is useful to note that altered expression of the gene in the hippocampus may be caused by the downregulation of some micro-RNAS, particularly of miR-185, found in the critical region deleted in 22q11.2 deletion syndrome (Forstner et al. 2013). Another target of miR-185 is *RHOA*, also altered in schizophrenia and involved in cortical interneuron migration, and is one of the genes showing strong signals of positive selection in AMHs compared to Neanderthals (Green et al. 2010). Two members of the CDC42 signaling pathway are also altered in schizophrenia: *CDC42EP4* (Datta et al. 2015), which is hypermethylated in AMHs compared to Denisovans (Gokhman et al. 2014), and *CDC42BPB* (Narayan et al. 2008), which is a target of FOXP2 (Spiteri et al. 2007). *ARHGAP32* is another partner of CDC42 related to schizophrenia and schizotypal personality traits (Ohi et al. 2012). It encodes a receptor of NMDA that modulates Rho-GTPase activity and it bears a fixed change (E1489D) in AMHs compared to Denisovans (Meyer et al. 2012). These data suggest that synergistic alterations in CDC42 signaling pathway may contribute to spine deficits in cells in schizophrenia and that this pathway has changed in our species. Concerning the SHH-GLI pathway, we expect it to have played a key role in the anatomical and physiological events leading to globularization (see Boeckx et al. submitted), but it also contributes to the pathobiology of schizophrenia (Boyd et al. 2015). SHH upregulates *DISC1* (Boyd et al. 2015). DISP1, one component of the SHH signalling network, shows a fixed change in AMHs (Green et al., 2010). SOX factors provide positional information in SHH-directed neural patterning together with GLI factors and some of them are related to schizophrenia. Hence, SOX10 is found to be hypermethylated in the brain of schizophrenics (Iwamoto et al. 2005, Wockner et al. 2014). Together with DISC1 it acts as negative regulator of oligodendrocyte differentiation (Drerup et al. 2009, Hattori et al. 2014). SOX2 is also involved in the enhancer effect of human endogenous retroviruses (HERVs) on brain genes related to schizophrenia, specifically on *PRODH* (Suntsova et al. 2013). Schizophrenia has been claimed to result in part from epigenetic changes that deregulate HERV-activity (Frank et al. 2005; Diem et al. 2012). HERVs are non-coding DNA remnants of retroviral infections occurred during primate evolution and seem to have fueled genomic rearrangements associated with or subsequent to speciation events (Böhne et al. 2008), so we expect them to have contributed as well to language evolution (see Benítez-Burraco and Uriagereka 2016 for discussion). Interestingly, a recent study by Castro-Nallar (2015) also found intriguing evidence of diversity in the schizophrenic oropharyngeal microbiome, with *Ascomycota* being more dominant and lactic acid being more abundant in schizophrenics than controls. The differences in bacteria between the two groups was clear, although its functional significance remains obscure. The microbiome has been shown to influence human cognition and behaviour through imbalances in the microbiota-gut-central nervous system axis (Foster and McVey Neufeld 2013, Hsiao et al. 2013). Other causal relations between schizophrenia and the ‘phageome’ have been posited (Yolken et al. 2015), and a number of studies connecting immune disorders and schizophrenia have also been forthcoming (reviewed by Severance et al. 2013). The behavioural and cognitive alterations seen in the microbiome can be changed via probiotic and antibiotic interventions (Jakobsson et al. 2010), and so an understanding of the relationship between cognition and viral, bacterial and fungal profiles could lead to successful remedial action. Together with Benítez-Burraco and Uriagereka’s (2016) claim that brain/immune system crosstalk led to alterations in brain connectivity giving rise to language, the microbiome appears to be a potentially fruitful area of research into the neurocognitive origins of schizophrenia.

On the whole, we believe that the genes reviewed in the last part of this section are important for both in the ethiopathology of schizophrenia and the evolution of language-readiness in the species, and provide a causative explanation to the origins and prevalence of schizophrenia. Importantly, the genes discussed here map onto specific neuronal types (mostly, GABAergic), particular brain areas (several cortical layers, thalamic nuclei), particular physiological processes (the balance between inhibition and excitation), specific developmental processes (inter and interhemispheric axon pathfinding), and particular cognitive abilities (formal thought), all of which are aspects known to be impaired in schizophrenia.

We wish to end by highlighting several similarities between the presumed Neanderthal head/brain/mind and the observed schizophrenia phenotype. The visual system changed in our species and this change surely had cognitive implications (discussed in Benítez-Burraco and Boeckx 2015). The changes involved not only orbit size reduction, but also anatomical and functional modifications at the level of the occipital lobe, which provides the roots of the visual system (Pearce et al. 2013), and of the frontal lobe (Masters et al. 2015). This entails that AMHs devote less of their brains to vision compared to Neanderthals, to the benefit of high-level cognitive processing (see Pearce et al. 2013 for details). A reduction of the visual area in AMHs probably led to an expansion of the parietal region in service of language (see Benítez-Burraco and Boeckx 2015 for detailed discussion). Interestingly, visual deficits either predispose towards or protect against schizophrenia for which language is crucially implicated (Silverstein et al. 2013; Leivada and Boeckx 2014). Schizophrenic patients exhibit impairments in visual information recall independent of working memory (Nuechterlein et al. 2004). Butler and Javitt (2005) claimed that visual-evoked potentials point to a selective impairment of the magnocellular pathway responsible for object motion and the interpretation of spatial relationships. Developments in optical coherence tomography (OCT) also permit more extensive study of the retina of schizophrenic patients (Schönfeldt-Lecuona et al. 2016). Given the oscillopathic perspective proposed here, it is significant that the impairments in Gestalt perception (Uhlhaas et al. 2006) and multimodal integration (Williams et al. 2010) documented in schizophrenia may be dependent on certain rhythm bands. Specifically, decreased *β* activity and altered *γ* phase coherence have been associated with poor Gestalt perception (Uhlhaas et al. 2006; Spencer et al. 2009), while weakened parietal *β*1 has been implicated in multimodal integration deficits (Kopell et al. 2011). Because each rhythm plays numerous, non-overlapping roles, it is crucial for these oscillopathic studies to be accompanied by biophysical modeling and computationally explicit mesoscopic frameworks of regionally localized cross-frequency coupling functionality. Adding to this, anomalies in eye development have been observed in schizophrenia (Leivada and Boeckx 2014). Moreover, some of the candidate genes for globularization play a role in eye development and are candidates for schizophrenia, like *SPAG5*, reviewed above (see Benítez-Burraco and Boeckx 2015 for details). These above considerations suggest that vision and visual cognition played a role as well in the emergence of language-readiness, but that, conversely, visual deficits in schizophrenia are causally linked to the cognitive dysfunction that is typical of the disorder.

## 5. Conclusions

The considerations we have made here may provide a suitable response to Dehaene et al.’s (2015: 2) observation that linguistic computation requires “a specific recursive neural code, as yet unidentified by electrophysiology, possibly unique to humans, and which may explain the singularity of human language and cognition”. Disruptions to the present dynomic model of linguistic computation may represent a comprehensive, unifying account of language-related neurocognitive disorders. While Crow (1997) famously argued that schizophrenia is the ‘price we paid for language’, we believe a more accurate claim is that schizophrenia is the price we paid for a globular braincase housing more efficient and widespread recursive oscillatory embeddings. This view should also, we believe, replace the still considerably strong grip psychoanalysis has on psychiatry, since the present hypotheses are broadly incompatible with the core psychoanalytic principle that symptoms are primarily the result of underlying traumas and emotional conflicts (see also Ceylan et al. 2016 for moves in this direction, with the authors claiming that neural synchronization explains ‘the psychoanalytic unconscious’). Hierarchical rhythmic coupling operations of the kind proposed in Murphy (2015a) and discussed here may also provide ways of integrating different forms of hierarchical representations, such as phonological, semantic and syntactic information (see Ding et al. 2016). As we have argued, schizophrenia is of particular interest because it represents a mode of cognition and externalization of thought distinct from, but plainly related to, normally functioning linguistic cognition. Importantly, this deviance seems construable in terms of an alteration of the cognome-dynome cross-talk. A dynomic perspective cuts across the traditional positive-negative symptom division, being implicated both in abnormal active processes and in the absence of normal functions.

Our view of schizophrenia as an oscillopathy is also in line with more general, recent moves in neuroscience to view psychiatric illnesses as oscillatory connectomopathies (Vinogradov and Herman 2016; Cao et al. 2016). The considerations we have presented also reinforce the view that the survey of abnormal cognitive/linguistic development in our species should help unravel the evolutionary itinerary followed by our faculty of language. The high number of candidates for schizophrenia selected in our species ostensibly proves this. At the same time, this history should help to understand the aetiology, clinical manifestations, and prevalence of schizophrenia. Significantly, the more novel a neural network is in evolutionary terms, the less resilient it is, due to its lack of robust compensatory mechanisms (Toro et al. 2010). Not surprisingly, brain development mirrors brain degeneration. Accordingly, the later a brain region develops, the earlier it degenerates in old age. This particularly holds for a transmodal network specifically associated with intellectual ability and episodic memory, which connects areas of increased vulnerability to schizophrenia (among other conditions) (Douaud et al. 2014). From an aetiological perspective, one plausible explanation for the high prevalence of schizophrenia (and other complex diseases) among human modern populations is that the same factors that prompted the transition from an ape-like cognition to a human-specific cognition (demographic bottlenecks, specific mutations, and cultural changes) uncovered cryptic variation and de-canalized primate cognition, which is composed of cognitive blocks which are particularly robust after millions of years of stabilizing selection (see Gibson 2009 for details). Plausibly, de-canalization explains as well why the number of disorders affecting human cognition is quite low: they may be the only possible phenotypes resulting from the interaction of the factors that regulate the development of the human brain and cognition. These possible phenotypes can be characterised as restricted areas within the adaptive landscape of the human cognitive phenotype (see McGhee 2006 for a general discussion on adaptive landscapes, and Benítez-Burraco 2016 for an account of language disorders from this evo-devo perspective). Finally, we further expect that the present proposal has the potential to provide robust endophenotypes of schizophrenia and contribute to an improved diagnosis and treatment of the disorder.

## Acknowledgements

Preparation of this work was supported in part by funds from the Spanish Ministry of Economy and Competitiveness (grant numbers FFI-2013-43823-P and FFI2014-61888-EXP to ABB) and in part by an Economic and Social Research Council scholarship (1474910).

## References

Aeby, A., De Tiège, X., Creuzil, M., David, P., Balériaux, D., Van Overmeire, B., et al. (2013). Language development at 2 years is correlated to brain microstructure in the left superior temporal gyrus at term equivalent age: a diffusion tensor imaging study. Neuroimage 78, 145–51.

Ainsworth, M., Lee, S., Cunningham, M.O., Roopun, A.K., Traub, R.D., Kopell, N.J. et al. (2011). Dual gamma rhythm generators control interlaminar synchrony in auditory cortex. J. Neurosci. 31: 17040–17051.

Anderson, G. R., Galfin, T., Xu, W., Aoto, J., Malenka, R. C., and Südhof, T. C. (2012). Candidate autism gene screen identifies critical role for cell-adhesion molecule CASPR2 in dendritic arborization and spine development. Proc. Natl. Acad. Sci. USA. 109, 18120–5.

Andreason, N.C., Hoffman, R.E., and Grove, W.M. (1985). “Mapping abnormalities in language and cognition”, in Controversies in Schizophrenia: Changes and Constancies. (New York: Guilford Press), 199–226.

Angrilli, A., Spironelli, C., Elbert, T., Crow, T.J., Marano, G., and Stegagnno, L. (2009). Schizophrenia as failure of left hemispheric dominance for the phonological component of language. PLoS ONE 4, e4507. doi:10.1371/journal.pone.0004507.

Anitha, A., Thanseem, I., Nakamura, K., Vasu, M. M., Yamada, K., Ueki, T., et al. (2014). Zinc finger protein 804A (ZNF804A) and verbal deficits in individuals with autism. J. Psychiatry Neurosci. 39, 294–303.

Arbib, M. A., Mundhenk, and T. N. (2005). Schizophrenia and the mirror system: an essay. Neuropsychologia 43, 268–80.

Bakhshi, K., and Chance, S. A. (2015). The neuropathology of schizophrenia: A selective review of past studies and emerging themes in brain structure and cytoarchitecture. Neuroscience 303, 82–102

Barr, M.S., Farzan, F., Rajji, T.K., Voineskos, A.N., Blumberger, D.M., Arenovich, T., et al. (2013). Can repetitive magnetic stimulation improve cognition in schizophrenia? Pilot data from a randomized controlled trial. Biol. Psychiatry 73, 510–517.

Basar-Eroǧlu, C., Mathes, B., Brand, A., and Schmiedt-Fehr, C. (2011). Occipital gamma response to auditory stimulation in patients with schizophrenia. Int. J. Psychophysiol. 79, 3–8.

Becker, J., Czamara, D., Hoffmann, P., Landerl, K., Blomert, L., Brandeis, D., et al. (2012). Evidence for the involvement of ZNF804A in cognitive processes of relevance to reading and spelling. Transl. Psychiatry 2, e136.

Benes, F. M., Lim, B., and Subburaju, S. (2009). Site-specific regulation of cell cycle and DNA repair in post-mitotic GABA cells in schizophrenic versus bipolars. Proc. Natl. Acad. Sci. U. S. A. 106, 11731–6.

Benes, F. M., Lim, B., Matzilevich, D., Walsh, J. P., Subburaju, S., Minns, and M. (2007). Regulation of the GABA cell phenotype in hippocampus of schizophrenics and bipolars. Proc. Natl. Acad. Sci. U.S.A. 104, 10164–9.

Benítez-Burraco, A. (2016). “A biolinguistic approach to language disorders: towards a paradigm shift in clinical linguistics”, in Advances in Biolinguistics: The Human Language Faculty and its Biological Basis (London: Routledge)

Benítez-Burraco, A., and Uriagereka, J. (2016). The immune syntax revisited: opening new windows on language evolution. Front. Mol. Neurosci. 8, 84. doi: 10.3389/fnmol.2015.00084.

Bennet, M. R. (2011). Schizophrenia: susceptibility genes, dendritic-spine pathology and gray matter loss. Prog. Neurobiol. 95, 275–300.

Benzel, I., Bansal, A., Browning, B. L., Galwey, N. W., Maycox, P. R., McGinnis, R., et al. (2007). Interactions among genes in the ErbB-Neuregulin signalling network are associated with increased susceptibility to schizophrenia. Behav. Brain Funct. 3, 31.

Bienvenu, T., Diebold, B., Chelly, J., and Isidor, B. (2013). Refining the phenotype associated with MEF2C point mutations. Neurogenetics 14, 71–5.

Blair, L. M., Pickler, R. H., and Anderson, C. (2015). Integrative review of genetic factors influencing neurodevelopmental outcomes in preterm infants. Biol. Res. Nurs.

Bleuler, E. (1911). Dementia Praecox or the Group of Schizophrenias. New York: International Universities Press.

Böhne, A., Brunet, F., Galiana-Arnoux, D.., Schultheis, C., and Volff, J. N. (2008). Transposable elements as drivers of genomic and biological diversity in vertebrates. Chromosome Res. 16: 203–15.

Booth, C. A., Brown, J. T., and Randall, A. D. (2014). Neurophysiological modification of CA1 pyramidal neurons in a transgenic mouse expressing a truncated form of disrupted-in-schizophrenia 1. Eur. J. Neurosci. 39, 1074–90.

Boyd, P. J., Cunliffe, V. T., Roy, S., and Wood, J. D. (2015). Sonic hedgehog functions upstream of disrupted-in-schizophrenia 1 (disc1): implications for mental illness. Biol. Open 4, 1336–43.

Bozza, M., Bernardini, L., Novelli, A., Brovedani, P., Moretti, E., Canapicchi, R., et al. (2013). 6p25 interstitial deletion in two dizygotic twins with gyral pattern anomaly and speech and language disorder. Eur. J. Paediatr. Neurol. 17, 225–31.

Braisted, J.E., Ringstedt, T., and O’Leary, D.D.M (2009). Slits are chemorepel-lents endogenous to hypothalamus and steer thalamocortical axons into ventral telencephalon. Cereb. Cortex 19, 144–151.

Brandon, N. J., and Sawa, A. (2011). Linking neurodevelopmental and synaptic theories of mental illness through DISC1. Nat. Rev. Neurosci. 12, 707–22.

Brüne, M. (2004). Schizophrenia-an evolutionary enigma? Neurosci. Biobehav Rev. 28, 41–53.

Buonanno, A. (2010). The neuregulin signaling pathway and schizophrenia: from genes to synapses and neural circuits. Brain Res. Bull. 83, 122–31.

Burdick, K. E., DeRosse, P., Kane, J. M., Lencz, T., Malhotra, A.K. (2010). Association of genetic variation in the MET proto-oncogene with schizophrenia and general cognitive ability. Am. J. Psychiatry 167, 436–43.

Butler, P.D., Javitt, D.C. (2005). Early-stage visual processing deficits in schizophrenia. Curr. Opin. Psychiatry 18, 151–157.

Cannon, T. D. (2015). How schizophrenia develops: cognitive and brain mechanisms underlying onset of psychosis. Trends Cogn. Sci. 19, 744–56.

Cao, H., Dixson, L., Meyer-Lindenberg, A., and Tost, H. (2016). Functional connectivity measures as schizophrenia intermediate phenotypes: advances, limitations, and future directions. Curr. Opin. Neurobiol. 36, 7–14.

Carvill, G. L., Regan, B. M., Yendle, S. C., O’Roak, B. J., Lozovaya, N., Bruneau, N., et al. (2013). GRIN2A mutations cause epilepsy-aphasia spectrum disorders. Nat. Genet. 45, 1073–6

Castro-Nallar, E., Bendall, M.L., Pérez-Losada, M., Sabuncyan, S., Severance, E.G., Dickerson, F.B., et al. (2015) Composition, taxonomy and functional diversity of the oropharynx microbiome in individuals with schizophrenia and controls. PeerJ 3, e1140. https://doi.org/10.7717/peerj.1140.

Ceylan, M.E., Dönmez, A., Ünsalver, B.O., and Evrensel, A. (2016). Neural synchronization as a hypothetical explanation of the psychoanalytic unconscious. Conscious. Cogn. 40, 34–44. http://dx.doi.org/10.1016/j.concog.2015.12.011.

Chiu, Y. C., Li, M. Y., Liu, Y. H., Ding, J. Y., Yu, J. Y., and Wang, T. W. (2014). Foxp2 regulates neuronal differentiation and neuronal subtype specification. Dev. Neurobiol. 74, 723–38.

Cho, K. K., Hoch, R., Lee, A. T., Patel, T., Rubenstein, J. L, and Sohal, V. S. (2015). Gamma rhythms link prefrontal interneuron dysfunction with cognitive inflexibility in Dlx5/6(+/-) mice. Neuron 85, 1332–43.

Chomsky, N. (1995). The Minimalist Program. Cambridge, MA: MIT Press.

Cohen, M. M. Jr. (2014). Proteus syndrome review: molecular, clinical, and pathologic features. Clin. Genet. 85, 111–9.

Courcet, J. B., Faivre, L., Malzac, P., Masurel-Paulet, A., López, E., Callier, P., et al. (2012). The DYRK1A gene is a cause of syndromic intellectual disability with severe microcephaly and epilepsy. J. Med. Genet. 49, 731–6.

Cousijn, H., Tunbridge, E. M., Rolinski, M., Wallis, G., Colclough, G. L., Woolrich, M. W., et al. (2015). Modulation of hippocampal theta and hippocampal-prefrontal cortex function by a schizophrenia risk gene. Hum. Brain Mapp. 36, 2387–95.

Crow, T. J. (1997). Is schizophrenia the price that Homo sapiens pays for language? Schizophr. Res. 28, 127–41.

Crow, T. J. (2008). The ‘big bang’ theory of the origin of psychosis and the faculty of language. Schizophr. Res. 102, 31–52.

Crow, T. J. (2013). The XY gene hypothesis of psychosis: origins and current status. Am. J. Med. Genet. B Neuropsychiatr. Genet. 162, 800–24.

Crow, T.J. (1980). Molecular pathology of schizophrenia: More than one disease process? Br. Med. J. 280, 66–68.

Crow, T.J. (1997). Is schizophrenia the price that Homo sapiens pays for language? Schizophr. Res. 28, 127–141. doi:10.1016/S0920-9964(97)00110-2.

Datta, D., Arion, D., Corradi, J. P., and Lewis, D. A. (2015). Altered expression of CDC42 signaling pathway components in cortical layer 3 pyramidal cells in schizophrenia. Biol. Psychiatry 78, 775–85.

De Ligt, J., Willemsen, M. H., van Bon, B. W. M., Kleefstra, T., Yntema, H. G., Kroes T, et al. (2012). Diagnostic exome sequencing in persons with severe intellectual disability. New Engl. J. Med. 367, 1921–1929.

Dehaene, S., Meyniel, F., Wacongne, C., Wang, L., and Pallier, C. (2015). The neural representation of sequences: from transition probabilities to algebraic patterns and linguistic trees. Neuron 88: 2–19.

Dennis, E. L., Jahanshad, N., Rudie, J. D., Brown, J. A., Johnson, K., McMahon, K. L., et al. (2011). Altered structural brain connectivity in healthy carriers of the autism risk gene, CNTNAP2. Brain Connect. 1, 447–59.

Diederen, K.M.J, De Weijer, A.D., Daalman, K., Blom, J.D., Neggers, S.F.W, Kahn, R.S., et al. (2010). Decreased language lateralization is characteristic of psychosis, not auditory hallucinations. Brain 133, 3734–3744. doi: 10.1093/brain/awq313.

Diem, O., Schäffner, M., Seifarth, W., and Leib-Mösch, C. (2012). Influence of antipsychotic drugs on human endogenous retrovirus (HERV) transcription in brain cells. PLoS One 7, e30054.

Dimassi, S., Labalme, A., Lesca, G., Rudolf, G., Bruneau, N., Hirsch, E., Arzimanoglou, A., et al. (2014). A subset of genomic alterations detected in rolandic epilepsies contains candidate or known epilepsy genes including GRIN2A and PRRT2. Epilepsia 55, 370–8.

Ding, N., Melloni, L., Zhang, H., Tian, X., and Poeppel, D. (2016). Cortical tracking of hierarchical linguistic structures in connected speech. Nat. Neurosci. 19, 158–64. doi:10.1038/nn.4186.

Djurovic, S., Gustafsson, O., Mattingsdal, M., Athanasiu, L., Bjella, T., Tesli, M., et al. (2010). A genome-wide association study of bipolar disorder in Norwegian individuals, followed by replication in Icelandic sample. J. Affect. Disord. 126, 312–6.

Domanico, D., Fragiotta, S., Trabucco, P., Nebbioso, M., and Vingolo, E. M. (2012). Genetic analysis for two italian siblings with usher syndrome and schizophrenia. Case Rep. Ophthalmol. Med. 2012, 380863.

Douaud, G., Groves, A. R., Tamnes, C. K., Westlye, L. T., Duff, E. P., Engvig, A., et al. (2014). A common brain network links development, aging, and vulnerability to disease. Proc. Natl. Acad. Sci USA. 111, 17648–53.

Drerup, C. M., Wiora, H. M., Topczewski, J., and Morris, J. A. (2009). Disc1 regulates foxd3 and sox10 expression, affecting neural crest migration and differentiation. Development 136, 2623–32.

Dudek, H., Datta, S. R., Franke, T. F., Birnbaum, M. J., Yao, R., Cooper, G. M., et al. (1997). Regulation of neuronal survival by the serine-threonine protein kinase Akt. Science 275(5300):661–5.

Egan, M. F., Goldberg, T. E., Kolachana, B. S., Callicott, J. H., Mazzanti, C. M., Straub, R. E., et al. (2001). Effect of COMT Val108/158 Met genotype on frontal lobe function and risk for schizophrenia. Proc. Natl. Acad. Sci. U.S.A. 98, 6917–6922.

Emamian, E. S., Karayiorgou, M., and Gogos, J. A. (2004). Decreased phosphorylation of NMDA receptor type 1 at serine 897 in brains of patients with schizophrenia. J. Neurosci. 24, 1561–4.

Endris, V., Wogatzky, B., Leimer, U., Bartsch, D., Zatyka, M., Latif, F., et al. (2002). The novel Rho-GTPase activating gene MEGAP/ srGAP3 has a putative role in severe mental retardation. Proc. Natl. Acad. Sci. U.S.A. 99, 11754–9.

Farzan, F., Barr, M.S., Sun, Y., Fitzgerald, P.B., Daskalakis, Z.J. (2012). Transcranial magnetic stimulation on the modulation of gamma oscillations in schizophrenia. Ann. N. Y. Acad. Sci. 1265, 25–35.

Ferrarelli, F., Massimini, M., Peterson, M.J., Riedner, B.A., Lazar, M., Murphy, M.J., et al. (2008). Reduced evoked gamma oscillations in the frontal cortex in schizophrenia patients: a TMS/EeG study. Am. J. Psychiatry 165, 996–1005.

Ferrarelli, F., Sarasso, S., Guller, Y., Riedner, B.A., Peterson, M.J., Bellesi, M., et al. (2012). Reduced natural oscillatory frequency of frontal thalamocortical circuits in schizophrenia. Arch. Gen. Psychiatry 69, 766–774. doi:10.1001/archgenpsychiatry.2012.147.

Fisahn, A., Neddens, J., Yan, L., and Buonanno, A. (2009). Neuregulin-1 modulates hippocampal gamma oscillations: implications for schizophrenia. Cereb. Cortex 19, 612–8.

Flint, J., and Munafò, M. R. (2014). Genetics: finding genes for schizophrenia. Curr. Biol. 24, R755–7.

Forstner, A. J., Degenhardt, F., Schratt, G., and Nöthen, M. M. (2013). MicroRNAs as the cause of schizophrenia in 22q11.2 deletion carriers, and possible implications for idiopathic disease: a mini-review. Front. Mol. Neurosci. 6, 47.

Foster, J.A., and McVey Neufeld, K-A. (2013). Gut-brain axis: how the microbiome influences anxiety and depression. Trends Neurosci. 36: 305–312.

Frank, O., Giehl, M., Zheng, C., Hehlmann, R., Leib-Mösch, C., and Seifarth, W. (2005). Human endogenous retrovirus expression profiles in samples from brains of patients with schizophrenia and bipolar disorders. J. Virol.79, 10890–901.

Fraser, W., King, K., Thomas, P., Kendall, R.E. (1986). The diagnosis of schizophrenia by language analysis. Br. J. Psychiatry 148, 275–278.

Frith, C.D. (1992). The Cognitive Neuropsychology of Schizophrenia. Hove: Lawrence Erlbaum Associates.

Frith, C.D., Allen, H.A. (1988). “Language disorders in schizophrenia and their implications for neuropsychology”, in Schizophrenia: The Major Issues (Oxford: Heinemann), 172–186.

Gattaz, W.F., Kohlmeyer, K., and Gasser, T. (1991). “Computer tomographic studies in schizophrenia”, in Search for the Causes of Schizophrenia. Vol. 2. Berlin: Springer.

Ghorashi, S., Spencer, K.M. (2015). Attentional load effects on beta oscillations in healthy and schizophrenic individuals. Front. Psychiatry 6, 149. doi: 10.3389/fpsyt.2015.00149.

Gibson, G. (2009). Decanalization and the origin of complex disease. Nature Rev. Genet. 10: 134–140.

Gilks, W. P., Hill, M., Gill, M., Donohoe, G., Corvin, A. P., and Morris, D. W. (2012). Functional investigation of a schizophrenia GWAS signal at the CDC42 gene. World J. Biol. Psychiatry 13, 550–4.

Gjørlund, M. D., Nielsen, J., Pankratova, S., Li, S., Korshunova, I., Bock, E., et al. (2012). Neuroligin-1 induces neurite outgrowth through interaction with neurexin-β and activation of fibroblast growth factor receptor-1. FASEB J. 26, 4174–86.

Godsil, B. P., Kiss, J. P., Spedding, M., and Jay, T. M. (2013). The hippocampal-prefrontal pathway: The weak link in psychiatric disorders? Eur. Neuropsychopharmacol. 23, 1165–1181.

Gonzalez-Burgos, G., Cho, R.Y., Lewis, D.A. (2015). Alterations in cortical network oscillations and parvalbumin neurons in schizophrenia. Biol. Psychiatry 77, 1031–1040.

Gracia-Sancho, J., Villarreal, G. Jr., Zhang, Y., and Garcia-Cardena, G. (2010). Activation of SIRT1 by resveratrol induces KLF2 expression conferring an endothelial vasoprotective phenotype. Cardiovasc. Res. 85, 514–519.

Grant, A., Fathalli, F., Rouleau, G., Joober, R., and Flores, C. (2012). Association between schizophrenia and genetic variation in DCC: a case-control study. Schizophr. Res. 137, 26–31.

Grant, A., Hoops, D., Labelle-Dumais, C., Prévost, M., Rajabi, H., Kolb, B., et al. (2007). Netrin-1 receptor-deficient mice show enhanced mesocortical dopamine transmission and blunted behavioural responses to amphetamine. Eur. J. Neurosci. 26, 3215–28.

Green, R. E., Krause, J., Briggs, A. W., Maricic, T., Stenzel, U., Kircher, M., et al. (2010). A draft sequence of the Neandertal genome. Science 328, 710–22.

Guilmatre, A., Huguet, G., Delorme, R., and Bourgeron, T. (2014). The emerging role of SHANK genes in neuropsychiatric disorders. Dev. Neurobiol. 74, 113–22.

Habib, M. R., Ganea, D. A., Katz, I. K., and Lamprecht, R. (2013). ABL1 in thalamus is associated with safety but not fear learning. Front. Syst. Neurosci. 7:5.

Hajós, M., Hoffmann, W. E., and Kocsis, B. (2008). Activation of cannabinoid-1 receptors disrupts sensory gating and neuronal oscillation: relevance to schizophrenia. Biol. Psychiatry 63, 1075–83.

Hall, J., Trent, S., Thomas, K. L., O’Donovan, M. C., and Owen, M. J. (2015). Genetic risk for schizophrenia: convergence on synaptic pathways involved in plasticity. Biol. Psychiatry 77, 52–8.

Hall, M. H., Taylor, G., Sham, P., Schulze, K., Rijsdijk, F., Picchioni, M., et al. (2011). The early auditory gamma-band response is heritable and a putative endophenotype of schizophrenia. Schizophr. Bull. 37, 778–87.

Hamdan, F. F., Daoud, H., Rochefort, D., Piton, A., Gauthier, J., Langlois, M., et al. (2010). De novo mutations in FOXP1 in cases with intellectual disability, autism, and language impairment. Am. J. Hum. Genet. 87, 671–678.

Hamdan, F. F., Srour, M., Capo-Chichi, J. M., Daoud, H., Nassif, C., Patry, L., et al. (2014). De novo mutations in moderate or severe intellectual disability. PLoS Genet. 10, e1004772.

Hämmerle, B., Carnicero, A., Elizalde, C., Ceron, J., Martínez, S., and Tejedor, F. J. (2003). Expression patterns and subcellular localization of the Down syndrome candidate protein MNB/DYRK1A suggest a role in late neuronal differentiation. Eur. J. Neurosci. 17, 2277–86.

Harrow, M., and Quinlan, D. (1977). Is disordered thinking unique to schizophrenia? Archives of General Psychiatry 28, 179–182.

Hashimoto, R., Ohi, K., Yasuda, Y., Fukumoto, M., Iwase, M., Iike, N., et al. (2010). The impact of a genome-wide supported psychosis variant in the ZNF804A gene on memory function in schizophrenia. Am. J. Med. Genet. B Neuropsychiatr. Genet. 153B, 1459–64

Hattori, T., Shimizu, S., Koyama, Y., Emoto, H., Matsumoto, Y., Kumamoto, N. et al. (2014). DISC1 (disrupted-in-schizophrenia-1) regulates differentiation of oligodendrocytes. PLoS One 9, e88506.

Hatzimanolis, A., McGrath, J. A., Wang, R., Li, T., Wong, PC., Nestadt, G., et al. (2013). Multiple variants aggregate in the neuregulin signaling pathway in a subset of schizophrenia patients. Transl. Psychiatry. 3, e264.

Henshall, K.R., Sergejew, A.A., Rance, G., McKay, C.M., Copolov, D.L. (2013). Interhemispheric EEG coherence is reduced in auditory cortical regions in schizophrenia patients with auditory hallucinations. Int. J. Psychophysiol. 89, 63–71.

Hinna, K. H., Rich, K., Fex-Svenningsen, Å., and Benedikz, E. (2015). The rat homolog of the schizophrenia susceptibility gene ZNF804A is highly expressed during brain development, particularly in growth cones. PLoS One 10, e0132456.

Hinzen, W., and Rosselló, J. (2015). The linguistics of schizophrenia: thought disturbance as language pathology across positive symptoms. Front. Psychol. 6, 971.

Hinzen, W., and Rosselló, J. (2015). The linguistics of schizophrenia: thought disturbance as language pathology across positive symptoms. Front. Psychol. 6, 971. doi:10.3389/fpsyg.2015.00971.

Hirayasu, Y., Shenton, M.E., Salisbury, D.F., Dickey, C.C., Fischer, I.A., Mazzoni, P., et al. (1998). Lower left temporal lobe MRI volumes in patients with first-episode schizophrenia compared with psychotic patients with first episode affective disorder and normal subjects. Am. J. Psychiatry 155, 1384–1391.

Hoffman, R.E., Rapaport, J., Mazure, C.M., Quinlan, D.M. (1999). Selective speech perception alterations in schizophrenic patients reporting hallucinated “voices”. Am. J. Psychiatry 156, 393–399.

Horn, H., Jann, K., Federspiel, A., Walther, S., Wiest, R., Müller, T., and Strik, W. (2012). Semantic network disconnection in formal thought disorder. Neuropsychobiology 66, 14–23.

Hou, X. J., Ni, K. M., Yang, J. M., and Li, X. M. (2014). Neuregulin 1/ErbB4 enhances synchronized oscillations of prefrontal cortex neurons via inhibitory synapses. Neuroscience. 261, 107–17.

Hou, X. J., Ni, K. M., Yang, J. M., and Li, X. M. (2014). Neuregulin 1/ErbB4 enhances synchronized oscillations of prefrontal cortex neurons via inhibitory synapses. Neuroscience 261, 107–17.

Hsiao, E.Y., McBride, S.W., Hsien, S., Sharon, G., Hyde, E.R., McCue, T., et al. (2013). Microbiota modulate behavioral and physiological abnormalities associated with neurodevelopmental disorders. Cell 155, 1451–1463.

Hui, L., Rao, W. W., Yu, Q., Kou, C., Wu, J. Q., He, J. C., et al. (2015). TCF4 gene polymorphism is associated with cognition in patients with schizophrenia and healthy controls. J. Psychiatr. Res. 69, 95–101.

Inda, M. C., DeFelipe, J., and Muñoz, A. (2006). Voltage-gated ion channels in the axon initial segment of human cortical pyramidal cells and their relationship with chandelier cells. Proc. Natl. Acad. Sci. USA. 103, 2920–5.

Inoue, K. I., Shiga, T., and Ito, Y. (2008). Runx transcription factors in neuronal development. Neural Dev. 3, 8104–3.

Ishii, R., Shinosaki, K., Ikejiri, I., Ukai, S., Yamashita, K., Iwase, M., et al. (2000). Theta rhythm increases in left superior temporal cortex during auditory hallucinations in schizophrenia: a case report. Neuroreport 11, 3283–3287.

Ivan Bon, B. W. M., Hoischen, A., Hehir-Kwa, J., de Brouwer, A. P. M., Ruivenkamp, C., Gijsbers, A. C. J., et al. (2011). Intragenic deletion in DYRK1A leads to mental retardation and primary microcephaly. Clin. Genet. 79: 296–99

Iwamoto, K., Bundo, M., Yamada, K., Takao, H., Iwayama-Shigeno, Y., Yoshikawa, T., et al. (2005). DNA methylation status of SOX10 correlates with its downregulation and oligodendrocyte dysfunction in schizophrenia. J. Neurosci. 25, 5376–81.

Jakobsson, H.E., Jernberg, C., Andersson, A.F., Sjölund-Karlsson, M., Jansson, J.K., and Engstrand, L. (2010). Short-term antibiotic treatment has differing long-term impacts on the human throat and gut microbiome. PLoS ONE 5, e9836.

Jenkins, A. K., Paterson, C., Wang, Y., Hyde, T. M., Kleinman, J. E., and Law, A. J. (2015). Neurexin 1 (NRXN1) splice isoform expression during human neocortical development and aging. Mol. Psychiatry doi: 10.1038/mp.2015.107.

Jensen, O., Gips, B., Bergmann, T.O., and Bonnefond, M. (2014). Temporal coding organized by coupled alpha and gamma oscillations prioritize visual processing. Trends Neurosci. 37, 357–369.

Jiang, L., Xu, Y., Zhu, X. T., Yang, Z., Li, H. J. Zuo, X. N. (2015). Local-to-remote cortical connectivity in early‐ and adulthood-onset schizophrenia. Transl. Psychiatry 5, e566.

Johnston, A., McBain, C. J., and Fisahn, A. (2014). 5-Hydroxytryptamine1A receptoractivation hyperpolarizes pyramidal cells and suppresses hippocampal gamma oscillations via Kir3 channel activation. J. Physiol. 592,4187–99.

Jung, J. S., Lee, H. J., Cho, C. H., Kang, S. G., Yoon, H. K., Park, Y. M., et al. (2014). Association between restless legs syndrome and CLOCK and NPAS2 gene polymorphisms in schizophrenia. Chronobiol. Int. 31, 838–44.

Kaczmarek, B.L.J (1987). “Regulatory function of the frontal lobes: a neurolinguistic perspective”, in The Frontal Lobes Revisited (New York: IRBN Press), 225–240.

Kamiya, A., Tan, P. L., Kubo, K., Engelhard, C., Ishizuka, K., Kubo, A., et al. (2008). Recruitment of PCM1 to the centrosome by the cooperative action of DISC1 and BBS4: a candidate for psychiatric illnesses. Arch. Gen. Psychiatry 65, 996–1006.

Karam, C. S., Ballon, J. S., Bivens, N. M., Freyberg, Z., Girgis, R. R., Lizardi-Ortiz, J. E., et al. (2010). Signaling pathways in schizophrenia: emerging targets and therapeutic strategies. Trends Pharmacol. Sci. 31, 381–90.

Katayama, K., Imai, F., Campbell, K., Lang, R. A., Zheng, Y., and Yoshida Y. (2013). RhoA and Cdc42 are required in pre-migratory progenitors of the medial ganglionic eminence ventricular zone for proper cortical interneuron migration. Development. 140, 3139–45.

Kersten, F. F., van Wijk, E., Hetterschijt, L., Bauβ, K., Peters, T. A., Aslanyan, M. G., et al. (2012). The mitotic spindle protein SPAG5/Astrin connects to the Usher protein network postmitotically. Cilia 1, 2.

Kircher, T.T.J, Oh, T.M., Brammer, M.J., McGuire, P.K. (2005). Neural correlates of syntax production in schizophrenia. Br. J. Psychiatry 186, 209–214.

Kishi, T., Fukuo, Y., Kitajima, T., Okochi, T., Yamanouchi, Y., Kinoshita, Y., et al. (2011). SIRT1 gene, schizophrenia and bipolar disorder in the Japanese population: an association study. Genes Brain Behav. 10, 257–63.

Kocsis, B. (2012). Differential role of NR2A and NR2B subunits in N-methyl-D-aspartate receptor antagonist-induced aberrant cortical gamma oscillations. Biol. Psychiatry 71, 987–95.

Konopka, G., Friedrich, T., Davis-Turak, J., Winden, K., Oldham, M. C., Gao, F.,et al. (2012). Human-specific transcriptional networks in the brain. Neuron 75, 601–17.

Kopell, N., Whittington, M., and Kramer, M. (2011). Neuronal assembly dynamics in the beta1 frequency range permits short-term memory. Proc. Natl. Acad. Sci. USA. 108, 3779–3784.

Kopell, N.J., Gritton, H.J., Whittington, M.A., Kramer, M.A. (2014). Beyond the connectome: the dynome. Neuron 83, 1319–1328.

Korotkova, T., Fuchs, E.C., Ponomarenko, A., von Engelhardt, J., and Monyer, H. (2010). NMDA receptor ablation on parvalbumin-positive interneurons impairs hippocampal synchrony, spatial representations, and working memory. Neuron 68, 557–569.

Kraeplin, E. (1913). Dementia Praecox and Paraphrenia. Edinburgh: Livingstone.

Kriebel, M., Wuchter, J., Trinks, S., and Volkmer H. (2012). Neurofascin: a switch between neuronal plasticity and stability. Int. J. Biochem. Cell. Biol. 44, 694–7.

Kromkamp, M., Uylings, H. B., Smidt, M. P., Hellemons, A. J., Burbach, J. P., and Kahn, R. S. (2003). Decreased thalamic expression of the homeobox gene DLX1 in psychosis. Arch. Gen. Psychiatry 60, 869–74.

Krug, A., Nieratschker, V., Markov, V., Krach, S., Jansen, A., Zerres, K., et al. (2010). Effect of CACNA1C rs1006737 on neural correlates of verbal fluency in healthy individuals. Neuroimage 49, 1831–6.

Kumar, D., Dedic, N., Flachskamm, C., Voulé, S., Deussing, J. M., and Kimura, M. (2015). Cacna1c (Cav1.2) modulates electroencephalographic rhythm and rapid eye movement sleep recovery. Sleep 38, 1371–80.

Kuperberg, G. R. (2010). Language in schizophrenia Part 1: an Introduction. Lang. Linguist Compass 4, 576–589.

Leivada, E. Boeckx, C. (2014). Schizophrenia and cortical blindness: Protective effects and implications for language. Front. Hum. Neurosci. 8, 940.

Lesca, G., Rudolf, G., Bruneau, N., Lozovaya, N., Labalme, A., Boutry-Kryza, N., et al. (2013). GRIN2A mutations in acquired epileptic aphasia and related childhood focal epilepsies and encephalopathies with speech and language dysfunction. Nat. Genet. 45, 1061–6.

Levchenko, A., Davtian, S., Freylichman, O., and Zagrivnaya, M., Kostareva, A., and Malashichev, Y. (2015). Beta-catenin in schizophrenia: Possibly deleterious novel mutation. Psychiatry Res. 228, 843–8.

Lewis, D.A., Curley, A.A., Glausier, J.R., Volk, D.W. (2012). Cortical parvalbumin interneurons and cognitive dysfunction in schizophrenia. Trends Neurosci. 35, 57–67.

Li, S., Weidenfeld, J., and Morrisey, E. E. (2004). Transcriptional and DNA binding activity of the Foxp1/2/4 family is modulated by heterotypic and homotypic protein interactions. Mol. Cell. Biol. 24, 809–822.

Li, T., Ma, X., Hu, X., Wang, Y., Yan, C., Meng, H., et al. (2008). PRODH gene is associated with executive function in schizophrenic families. Am. J. Med. Genet. B Neuropsychiatr. Genet. 147B, 654–7.

Li, T., Zeng, Z., Zhao, Q., Wang, T., Huang, K., Li, J., et al. (2013). FoxP2 is significantly associated with schizophrenia and major depression in the Chinese Han population. World J. Biol. Psychiatry 14, 146–50.

Li, W., Guo, X., and Xiao, S. (2015). Evaluating the relationship between reelin gene variants (rs7341475 and rs262355) and schizophrenia: A meta-analysis. Neurosci. Lett. 609, 42–7.

Li, X., Alapati, V., Jackson, C., Xia, S., Bertisch, H. C., Branch, C. A., et al. (2012). Structural abnormalities in language circuits in genetic high-risk subjects and schizophrenia patients. Psychiatry Res. 201, 182–9.

Li, X., Branch, C. A., and DeLisi, L. E. (2009). Language pathway abnormalities in schizophrenia: a review of fMRI and other imaging studies. Curr. Opin. Psychiatry 22, 131–9.

Linden, D. E., Lancaster, T. M., Wolf, C., Baird, A., Jackson, M. C., Johnston, S. J., et al. (2012). ZNF804A genotype modulates neural activity during working memory for faces. Neuropsychobiology 67, 84–92.

Linkenkaer-Hansen, K., Smit, D. J., Barkil, A., van Beijsterveldt, T. E., Brussaard, A. B., Boomsma, D. I., et al. (2007). Genetic contributions to long-range temporal correlations in ongoing oscillations. J. Neurosci. 27, 13882–9.

Liu, X., Somel, M., Tang, L., Yan, Z., Jiang, X., Guo, S. et al.(2012). Extension of cortical synaptic development distinguishes humans from chimpanzees and macaques. Genome Res.22, 611–22.

Ma, Y., Mi, Y.-J., Dai, Y.-K., Fu, H.-L., Cui, D.-X. and Jin, W. L. (2013). The inverse F-BAR domain protein srGAP2 acts through srGAP3 to modulate neuronal differentiation and neurite outgrowth of mouse neuroblastoma cells. PLoS One 8, e57865.

Malhotra, A. K., Kestler, L. J., Mazzanti, C., Bates, J. A., Goldberg, T., and Goldman, D. (2002). A functional polymorphism in the COMT gene and performance on a test of prefrontal cognition. Am. J. Psychiatry 159, 652–654.

Manoach, D. S., Pan, J. Q., Purcell, S. M., and Stickgold, R. (2015). Reduced sleep spindles in schizophrenia: a treatable endophenotype that links risk genes to impaired cognition? Biol. Psychiatry pii: S0006-3223(15)00818–5. doi: 10.1016/j.biopsych.2015.10.003.

Masters, M., Bruner, E., Queer, S., Traynor, S., and Senjem, J. (2015). Analysis of the volumetric relationship among human ocular, orbital and fronto-occipital cortical morphology. J. Anat. 227, 460–73.

Matsunoshita, Y., Ijiri, K., Ishidou, Y., Nagano, S., Yamamoto, T., Nagao, H., et al. (2011). Suppression of osteosarcoma cell invasion by chemotherapy is mediated by urokinase plasminogen activator activity via up-regulation of EGR1. PLoS One 6, e16234.

McCarthy, R.A., Warrington, E.K. (1990). Cognitive Neuropsychology. London: Academic Press.

McCarthy, S. E., McCombie, W. R., and Corvin, A. (2014). Unlocking the treasure trove: from genes to schizophrenia biology. Schizophr. Bull. 40, 492–6.

McClain, L. (1983). Encoding and retrieval in schizophrenics’ free recall. J. Nerv. Ment. Dis. 171, 471–479.

McGhee, G. R. (2006). The Geometry of Evolution: Adaptive Landscapes and Theo-retical Morphospaces. Cambridge: Cambridge University Press.

McKenna P.J., Oh, T.M. (2005). Schizophrenic Speech: Making Sense of Bathroots and Ponds that Fall in Doorways. Cambridge: Cambridge University Press.

Meyer, M., Kircher, M., Gansauge, M. T., Li, H., Racimo, F., Mallick, S., et al. (2012). A high-coverage genome sequence from an archaic Denisovan individual. Science 338, 222–6.

Minzenberg, M.J., Laird, A.R., Thelen, S., Carter, C.S., Glahn, D.C. (2009). Meta-analysis of 41 functional neuroimaging studies of executive function in schizophrenia. Arch. Gen. Psychiatry 66, 811–822.

Mitchell, A. C., Jiang, Y., Peter, C., and Akbarian, S. (2015). Transcriptional regulation of GAD1 GABA synthesis gene in the prefrontal cortex of subjects with schizophrenia. Schizophr. Res. 167, 28–34.

Molina, V., Papiol, S., Sanz, J., Rosa, A., Arias, B., Fatjó-Vilas, M., et al. (2011). Convergent evidence of the contribution of TP53 genetic variation (Pro72Arg) to metabolic activity and white matter volume in the frontal lobe in schizophrenia patients. Neuroimage 56, 45–51.

Moran, L. V., and Hong, L. E. (2011). High vs low frequency neural oscillations in schizophrenia. Schizophr. Bull. 37, 659–63.

Moran, L.V., Hong, L.E. (2011). High versus low frequency neural oscillations in schizophrenia. Schizophr. Bull. 37, 659–663.

Morice, R., and Ingram, J.C.L. (1982). Language analysis in schizophrenia. Aust. N. Z. J. Psychiatry 16, 11–21. doi:10.3109/00048678209161186.

Morice, R., and McNicol, D. (1985). The comprehension and production of complex syntax in schizophrenia. Cortex 21, 567–80.

Morice, R., and McNicol, D. (1986). Language changes in schizophrenia. Schizophr. Bull. 12, 239–250. doi:10.1093/schbul/12.2.239.

Mukai, J., Liu, H., Burt, R. A., Swor, D. E., Lai, W. S., Karayiorgou, M. et al. (2004). Evidence that the gene encoding ZDHHC8 contributes to the risk of schizophrenia. Nat. Genet. 36: 725–731.

Mukamel, Z., Konopka, G., Wexler, E., Osborn, G. E., Dong, H., Bergman, M. Y., et al. (2011). Regulation of MET by FOXP2, genes implicated in higher cognitive dysfunction and autism risk. J. Neurosci. 31, 11437–42.

Murphy, E. (2015a). The brain dynamics of linguistic computation. Front. Psychol. 6, 1515. doi: 10.3389/fpsyg.2015.01515.

Murphy, E. (2015b). Reference, phases and individuation: Topics at the labeling-interpretive interface. Opticon1826 17, 1–13. doi:10.5334/opt.cn.

Murphy, E. (2015c). Labels, cognomes and cyclic computation: An ethological perspective. Front. Psychol. 6, 715. doi: 10.3389/fpsyg.2015.00715.

Nakamura, T., Matsumoto, J., Takamura, Y., Ishii, Y., Sasahara, M., Ono, T, et al. (2015). Relationships among parvalbumin-immunoreactive neuron density, phase-locked gamma oscillations, and autistic/schizophrenic symptoms in PDGFR-β knock-out and control mice. PLoS One 10, e0119258.

Narayan, S., Tang, B. Head, S. R., Gilmartin, T. J., Sutcliffe, J. G., Dean, B., et al. (2008). Molecular profiles of schizophrenia in the CNS at different stages of illness. Brain Res. 1239, 235–48.

Narita, H. (2014). Endocentric Structuring of Projection-free Syntax. Amsterdam: John Benjamins.

Newbury, D. F., Paracchini, S., Scerri, T. S., Winchester, L., Addis, L., Richardson, A. J., et al. (2011). Investigation of dyslexia and SLI risk variants in reading‐ and language-impaired subjects. Behav. Genet. 41, 90–104.

Nguyen, P. T., Nakamura, T., Hori, E., Urakawa, S., Uwano, T., Zhao, J., et al. (2011). Cognitive and socio-emotional deficits in platelet-derived growth factor receptor-β gene knockout mice. PLoS One 6, e18004.

Ni, X., Trakalo, J., Valente, J., Azevedo, M. H., Pato, M. T., Pato, C. N., et al. (2005). Human p53 tumor suppressor gene (TP53) and schizophrenia: case-control and family studies. Neurosci. Lett. 388, 173–8.

Nicodemus, K. K., Elvevåg, B., Foltz, P. W., Rosenstein, M., Diaz-Asper, C., and Weinberger, D. R. (2014). Category fluency, latent semantic analysis and schizophrenia: a candidate gene approach. Cortex 55, 182–91.

Nuechterlein, K.H., Barch, D.M., Gold, J.M., Goldberg, T.E., Green, M.F., Heaton, R.K. (2004). Identification of separable cognitive factors in schizophrenia. Schizophr. Res. 72, 2–39.

Nunez, P.L., and Srinivasan, R. (2006). Electric Fields of the Brain: The Neurophysics of EEG. 2nd ed. Oxford: Oxford University Press.

Nunez, P.L., Wingeier, B.M., Silberstein, R.B. (2001). Spatial-temporal structures of human alpha rhythms: theory, microcurrent sources, multiscale measurements, and global binding of local networks. Hum. Brain Mapp. 13, 125–164.

Ogawa, L. M., and Vallender, E. J. (2014). Evolutionary conservation in genes underlying human psychiatric disorders. Front. Hum. Neurosci. 8, 283.

Ohi, K., Hashimoto, R., Nakazawa, T., Okada, T., Yasuda, Y., Yamamori, H., et al. (2012). The p250GAP gene is associated with risk for schizophrenia and schizotypal personality traits. PLoS One 7, e35696.

Oksenberg, N., and Ahituv, N. (2013). The role of AUTS2 in neurodevelopment and human evolution. Trends Genet. 29, 600–8.

Oksenberg, N., Stevison, L., Wall, J. D., and Ahituv, N. (2013). Function and regulation of AUTS2, a gene implicated in autism and human evolution. PLoS Genet. 9, e1003221

Onwuameze, O. E., Nam, K. W., Epping, E. A., Wassink, T. H., Ziebell, S., Andreasen, N. C., et al. (2013). MAPK14 and CNR1 gene variant interactions: effects on brain volume deficits in schizophrenia patients with marijuana misuse. Psychol. Med. 43, 619–31.

Oribe, N., Onitsuka, T., Hirano, S., Hirano, Y., Maekawa, T., Obayashi, C., et al. (2010). Differentiation between bipolar disorder and schizophrenia revealed by neural oscillation to speech sounds: an MEG study. Bipolar Disord. 12, 804–812. doi: 10.1111/j.1399-5618.2010.00876.x.

Osbun, N., Li, J., O’Driscoll, M. C., Strominger, Z., Wakahiro, M., Rider, E., et al. (2011). Genetic and functional analyses identify DISC1 as a novel callosal agenesis candidate gene. Am. J. Med. Genet. A. 155A, 1865–76.

O’Tuathaigh, C. M., Desbonnet, L., Moran, P. M., and Waddington, J. L. (2012). Susceptibility genes for schizophrenia: mutant models, endophenotypes and psychobiology. Curr. Top. Behav. Neurosci. 12, 209–50.

Ousley, O., Rockers, K., Dell, M. L., Coleman, K., and Cubells, J. F. (2007). A review of neurocognitive and behavioral profiles associated with 22q11 deletion syndrome: implications for clinical evaluation and treatment. Curr. Psychiatry Rep. 9, 148–58.

Owen, F., Cross, A.J., Crow, T.J., Longden, A., Poulter, M., Riley, G.J. (1978). Increased dopamine-receptor sensitivity in schizophrenia. Lancet 312, 223–226.

Pääbo, S. (2014). The human condition-a molecular approach. Cell 157, 216–26.

Papaleo, F., Lipska, B. K., and Weinberger, D. R. (2012). Mouse models of genetic effects on cognition: relevance to schizophrenia. Neuropharmacology 62, 1204–20.

Paskulin, D. d., Paixâo-Côrtes, V. R., Hainaut, P., Bortolini, M. C., and Ashton-Prolla, P. (2012). The TP53 fertility network. Genet. Mol. Biol. 35, 939–46.

Pearce, E., Stringer, C., and Dunbar, R. I. M. (2013). New insights into differences in brain organization between Neanderthals and anatomically modern humans. Proc. Biol. Sci. 280, 20130168.

Pearlson, G. D., and Folley, B. S. (2008). Schizophrenia, psychiatric genetics, and Darwinian psychiatry: an evolutionary framework. Schizophr. Bull. 34, 722–33.

Peng, X. D., Xu, P. Z., Chen, M. L., Hahn-Windgassen, A., Skeen, J., and Jacobs, J. (2003). Dwarfism, impaired skin development, skeletal muscle atrophy, delayed bone development, and impeded adipogenesis in mice lacking Akt1 and Akt2. Genes Dev. 17, 1352–1365.

Pérez-Santiago, J., Diez-Alarcia, R., Callado, L. F., Zhang, J. X., Chana, G., White, C. H., et al. (2012). A combined analysis of microarray gene expression studies of the human prefrontal cortex identifies genes implicated in schizophrenia. J. Psychiatr. Res. 46, 1464–74.

Peter, B., Raskind, W. H., Matsushita, M., Lisowski, M., Vu, T., Berninger, V. W., et al. (2011) Replication of CNTNAP2 association with nonword repetition and support for FOXP2 association with timed reading and motor activities in a dyslexia family sample. J. Neurodev. Disord. 3, 39–49.

Pittman-Polletta, B. R., Kocsis, B., Vijayan, S., Whittington, M. A., and Kopell, N. J. (2015). Brain rhythms connect impaired inhibition to altered cognition in schizophrenia. Biol. Psychiatry 77, 1020–30.

Pleasure, S. J., Anderson, S., Hevner, R., Bagri, A., Marin, O., Lowenstein, D. H., et al. (2000). Cell migration from the ganglionic eminences is required for the development of hippocampal GABAergic interneurons. Neuron 28, 727–40.

Plum, F. (1972). Prospects for research on schizophrenia. 3. Neurophysiology. Neuropathological findings. Neurosci. Res. Program Bull. 10, 384–388.

Poot, M. (2015). Connecting the CNTNAP2 networks with neurodevelopmental disorders. Mol. Syndromol. 6, 7–22.

Poot, M., van’t Slot, R., Leupert, R., Beyer, V., Passarge, E., and Haaf, T. (2009). Three de novo losses and one insertion within a pericentric inversion of chromosome 6 in a patient with complete absence of expressive speech and reduced pain perception. Med. Genet. 52, 27–30.

Popov, T., and Popova, P. (2015). Same clock, different time read-out: spontaneous brain oscillations and their relationship to deficient coding of cognitive content. NeuroImage 119, 316–324. http://dx.doi.org/10.1016/j.neuroimage.2015.06.071.

Potkin, S. G., Macciardi, F., Guffanti, G., Fallon, J. H., Wang, Q., Turner, J. A., et al. (2010). Identifying gene regulatory networks in schizophrenia. Neuroimage 53, 839–47.

Prüfer, K., Racimo, F., Patterson, N., Jay, F., Sankararaman, S., Sawyer, S., et al. (2014). The complete genome sequence of a Neanderthal from the Altai Mountains. Nature. 505(7481):43–9.

Ramsden, P. (2013). Understanding Abnormal Psychology: Clinical and Biological Perspectives. London: SAGE Publications.

Roberts, J. L., Hovanes, K., Dasouki, M., Manzardo, A. M., and Butler, M. G. (2014). Chromosomal microarray analysis of consecutive individuals with autism spectrum disorders or learning disability presenting for genetic services. Gene 535, 70–8.

Rockland, K. S. (2015). About connections. Front. Neuroanat. 9, 61.

Roll, P., Rudolf, G., Pereira, S., Royer, B., Scheffer, I. E., Massacrier, A., et al. (2006). SRPX2 mutations in disorders of language cortex and cognition. Hum. Mol. Genet. 15, 1195–207.

Roll, P., Vernes, S. C., Bruneau, N., Cillario, J., Ponsole-Lenfant, M., Massacrier, A., et al. (2010). Molecular networks implicated in speech-related disorders: FOXP2 regulates the SRPX2/uPAR complex. Hum. Mol. Genet. 19, 4848–60.

Rolstad, S., Pålsson, E., Ekman, C. J., Eriksson, E., Sellgren, C., and Landén, M. (2015). Polymorphisms of dopamine pathway genes NRG1 and LMX1A are associated with cognitive performance in bipolar disorder. Bipolar Disord. 17, 859–68.

Roux, F., Wibral, M., Mohr, H.M., Singer, W., Uhlhaas, P.J. (2012). Gamma-band activity in human prefrontal cortex codes for the number of relevant items maintained in working memory. J. Neurosci. 32, 12411–12420.

Royer-Zemmour, B., Ponsole-Lenfant, M., Gara, H., Roll, P., Lévêque, C., Massacrier, A., et al. (2008). Epileptic and developmental disorders of the speech cortex: ligand/receptor interaction of wild-type and mutant SRPX2 with the plasminogen activator receptor uPAR. Hum. Mol. Genet. 17, 3617–30.

Ruzicka, W. B., Subburaju, S., and Benes, F. M. (2015). Circuit‐ and diagnosis-specific DNA methylation changes at γ-aminobutyric acid-related genes in postmortem human hippocampus in schizophrenia and bipolar disorder. JAMA Psychiatry 72, 541–51.

Sans-Sansa, B., McKenna, P.J., Canales-Rodríguez, E.J., Ortiz-Gil, K., López-Araquistain, L., Sarró, S., et al. (2013). Association of formal thought disorder in schizophrenia with structural brain abnormalities in language-related cortical regions. Schizophr. Res. 146, 308–313. doi:10.1016/j.schres.2013.02.032.

Sartorius, N., Jablensky, A., Korten, A., Ernberg, G., Anker, M., Cooper, J.E., et al. (1986). Early manifestations and first-contact incidence of schizophrenia in different cultures. A preliminary report on the initial evaluation phase of the WHO Collaborative Study on determinants of outcomes of severe mental disorders. Psychol. Med. 16, 909–928.

Schönfeldt-Lecuona, C., Kregel, T., Schmidt, A., Pinkhardt, E.H., Lauda, F., Kassubek, J., et al. (2016). From imaging the brain to imaging the retina: optical coherence tomography (OCT) in schizophrenia. Schizophr. Bull. 42, 9–14. doi: 10.1093/schbul/sbv073.

Schulman, J.J., Cancro, R., Lowe, S., Lu, F., Walton, K.D., and Llinâs, R.R. (2011). Imaging of thalamocortical dysrhythmia in neuropsychiatry. Front. Hum. Neuroscience 5, 69. doi: 10.3389/fnhum.2011.00069.

Scott-Van Zeeland, A. A., Abrahams, B. S., Alvarez-Retuerto, A. I., Sonnenblick, L. I., Rudie, J. D., Ghahremani, D., et al. (2010). Altered functional connectivity in frontal lobe circuits is associated with variation in the autism risk gene CNTNAP2. Sci Transl Med. 2:56—80.

Seshadri, S., Faust, T., Ishizuka, K., Delevich, K., Chung, Y., Kim, S. H., et al. (2015). Interneuronal DISC1 regulates NRG1-ErbB4 signalling and excitatory-inhibitory synapse formation in the mature cortex. Nat. Commun. 6, 10118.

Severance, E.G., Gressitt, K.L., Stallings, C.R., Origoni, A.E., Khushalani, S., Leweke, F.M., et al. (2013). Discordant patterns of bacterial translocation markers and implications for innate immune imbalances in schizophrenia. Schizophr. Res. 148, 130–137.

Shulha, H. P., Crisci, J. L., Reshetov, D., Tushir, J. S., Cheung, I., Bharadwaj, R., et al (2012). Human-specific histone methylation signatures at transcription start sites in prefrontal neurons. PLoS Biol. 10, e1001427.

Silverstein, S. M., Wang, Y., and Keane, B. P. (2013). Cognitive and neuroplasticity mechanisms by which congenital or early blindness may confer a protective effect against. Front. Psychol. 3, 624.

Sollis, E., Graham, S. A., Vino, A., Froehlich, H., Vreeburg, M., Dimitropoulou, D., et al. (2015). Identification and functional characterization of de novo FOXP1 variants provides novel insights into the etiology of neurodevelopmental disorder. Hum. Mol. Genet. pii: ddv495.

Somel M, Liu X, and Khaitovich P. (2013). Human brain evolution: transcripts, metabolites and their regulators. Nat. Rev. Neurosci. 14, 112–127.

Souchet, B., Guedj, F., Sahøn, I., Duchon, A., Daubigney, F., Badel, A., et al. (2014). Excitation/inhibition balance and learning are modified by Dyrk1a gene dosage. Neurobiol. Dis. 69, 65–75.

Španiel, F., Horâcek, J., Tintěra, J., Ibrahim, I., Novâk, T., Cermâk, J., et al. (2011). Genetic variation in FOXP2 alters grey matter concentrations in schizophrenia patients. Neurosci. Lett. 493, 131–5.

Speevak, M. D. Farrell, S. A. (2011). Non-syndromic language delay in a child with disruption in the Protocadherin11X/Y gene pair. Am. J. Med. Genet. B Neuropsychiatr. Genet. 156, 484–9.

Spencer, K.M., Niznikiewicz, M.A., Nestor, P.G., Shenton, M.E., McCarley, R.W. (2009). Left auditory cortex gamma synchronization and auditory hallucination symptoms in schizophrenia. BMC Neurosci. 10, 85.

Spironelli, C., and Angrilli, A. (2015). Language-related gamma EEG frontal reduction is associated with positive symptoms in schizophrenia patients. Schizophr. Res. 165, 22–29. http://dx.doi.org/10.1016/j.schres.2015.04.003.

Spironelli, C., and Angrilli, A., Calogero, A., and Stegagno, L. (2011). Delta EEG band as a marker of left hypofrontality for language schizophrenia patients. Schizophr. Bull. 37, 757–767. doi:10.1093/schbul/sbp145.

Spiteri, E., Konopka, G., Coppola, G., Bomar, J., Oldham, M., Ou, J., et al. (2007). Identification of the transcriptional targets of FOXP2, a gene linked to speech and language, in developing human brain. Am. J. Hum. Genet. 81, 1144–57.

Srinivasan, S., Bettella, F., Mattingsdal, M., Wang, Y., Witoelar, A., Schork, A. J., et al. (2015). Genetic markers of human evolution are enriched in schizophrenia. Biol. Psychiatry, S0006-3223(15)00855–0.

Stein, C. M., Schick, J. H., Gerry Taylor, H., Shriberg, L. D., Millard, C., Kundtz-Kluge, A., et al. (2004). Pleiotropic effects of a chromosome 3 locus on speech-sound disorder and reading. Am. J. Hum. Genet. 74, 283–97.

Stephane, M., Kuskowski, M., and Gundel, J. (2014). Abnormal dynamics of language in schizophrenia. Psychiatry Res. 216, 320–4

Stephane, M., Pellizzer, G., Fletcher, C. R., and McClannahan, K. (2007). Empirical evaluation of language disorder in schizophrenia. J. Psychiatry Neurosci. 32, 250–8.

Stephens, A. S., and Morrison, N. A. (2014). Novel target genes of RUNX2 transcription factor and 1,25-dihydroxyvitamin D3. J. Cell. Biochem. 115, 1594–608.

Südhof, T. C. (2008). Neuroligins and neurexins link synaptic function to cognitive disease. Nature 455, 903–911.

Sun, J., Tang, Y., Lim, K.O., Wang, J., Tong, S., Li, H., et al. (2014). Abnormal dynamics of EEG oscillations in schizophrenia patients on multiple time scales. IEEE Trans. Biomed. Eng. 61, 1756–1764.

Suntsova, M., Gogvadze, E. V., Salozhin, S., Gaifullin, N., Eroshkin, F., Dmitriev, S. E., et al. Human-specific endogenous retroviral insert serves as an enhancer for the schizophrenia-linked gene PRODH. Proc. Natl. Acad. Sci. U.S.A. 110, 19472–7.

Szklarczyk, D., Franceschini, A., Wyder, S., Forslund, K., Heller, D., Huerta-Cepas, J., et al. (2015). STRING v10: protein-protein interaction networks, integrated over the tree of life. Nucleic Acids Res. 43, D447–52.

Tamlyn, D., McKenna, P.J., Mortimer, A.M., Lund, C.E., Hammond, S., Baddeley, A.D. (1992). Memory impairment in schizophrenia: its extent, affiliations and neuropsychological character. Psychol. Med. 22, 101–115. doi: 10.1017/S0033291700032773.

Tan, G. C., Doke, T. F., Ashburner, J., Wood, N. W., and Frackowiak, R. S. (2010). Normal variation in fronto-occipital circuitry and cerebellar structure with an autism-associated polymorphism of CNTNAP2. Neuroimage 53, 1030–42.

Teffer, K., and Semendeferi, K. (2012). Human prefrontal cortex: evolution, development, and pathology. Prog. Brain Res. 195, 191–218.

Thomas, P., King, K., Fraser, W., Kendall, R.E. (1987). Linguistic performance in schizophrenia: a comparison of patients with positive and negative symptoms. Acta Psychiatr. Scand. 76, 1-4–4-151.

Thygesen, J. H., Zambach, S. K., Ingason, A., Lundin, P., Hansen, T., Bertalan, M. et al. (2015). Linkage and whole genome sequencing identify a locus on 6q25-26 for formal thought disorder and implicate MEF2A regulation. Schizophr. Res. 169, 441–6.

Tosato, S., Bellani, M., Bonetto, C., Ruggeri, M., Perlini, C., Lasalvia, A., et al. Is neuregulin 1 involved in determining cerebral volumes in schizophrenia? Preliminary results showing a decrease in superior temporal gyrus volume. Neuropsychobiology 65, 119–25.

Turner, S. J., Mayes, A. K., Verhoeven, A., Mandelstam, S. A., Morgan, A. T., and Scheffer, I. E. (2015). GRIN2A: an aptly named gene for speech dysfunction. Neurology 84, 586–93.

Uhlhaas, P. J., Haenschel, C., Nikolic, D., and Singer, W. (2008). The role of oscillations and synchrony in cortical networks and their putative relevance for the pathophysiology of schizophrenia. Schizophr Bull. 34, 927–943.

Uhlhaas, P.J., and Singer, W. (2015). Oscillations and neuronal dynamics in schizophrenia: the search for basic symptoms and translational opportunities. Biol. Psychiatry 77, 1001–1009. http://dx.doi.org/10.1016/j.biopsych.2014.11.019.

Uhlhaas, P.J., Haenschel, C., Nikolic, D., and Singer, W. (2008). The role of oscillations and synchrony in cortical networks and their putative relevance for the pathophysiology of schizophrenia. Schizophr. Bull. 34, 927–943.

Uhlhaas, P.J., Linden, D.E., Singer, W., Haenschel, C., Lindner, M., Maurer, K., et al. (2006). Dysfunctional long-range coordination of neural activity during gestalt perception in schizophrenia. J. Neurosci. 26, 8168–8175.

van Bon, B. W., Balciuniene, J., Fruhman, G., Nagamani, S. C., Broome, D. L., Cameron, E., et al. (2011). The. phenotype. of. recurrent. 10q22q23 deletions and duplications. Eur. J. Hum. Genet. 19, 400–8.

van Os, J., and Kapur, S. (2009). Schizophrenia. Lancet 374, 635–45.

Verbrugghe, P., Bouwer, S., Wiltshire, S., Carter, K., Chandler, D., Cooper, M., et al. (2012). Impact of the Reelin signaling cascade (ligands-receptors-adaptor complex) on cognition in schizophrenia. Am. J. Med. Genet B Neuropsychiatr. Genet. 159B, 392–404.

Vernes, S. C., Newbury, D. F., Abrahams, B. S., Winchester, L., Nicod, J., Groszer, M., et al. (2008). A functional genetic link between distinct developmental language disorders. N. Engl. J. Med. 359, 2337–45.

Veyrac, A., Besnard, A., Caboche, J., Davis, S., and Laroche S. (2014). The transcription factor Zif268/Egr1, brain plasticity, and memory. Prog. Mol. Biol. Transl. Sci. 122, 89–129.

Vinogradov, S., and Herman, A. (2016). Psychiatric illnesses as oscillatory connectomopathies. Neuropsychopharmacology 41, 387–388. doi:10.1038/npp.2015.308.

Walker, R. M., Hill, A. E., Newman, A. C., Hamilton, G., Torrance, H. S., Anderson, S. M., et al. (2012). The DISC1 promoter: characterization and regulation by FOXP2. Hum. Mol. Genet. 21, 2862–72.

Waltereit, R., Leimer, U., von Bohlen und Halbach, O., Panke, J., Holter, S. M., Garrett, L., et al. (2012). Srgap37“ mice present a neurodevelopmental disorder with schizophrenia-related intermediate phenotypes. FASEB J. 26, 4418–28.

Waltereit, R., von Bohlen Leimer, U., und Halbach, O., Panke, J., Holter, S. M., Garrett, L., et al. (2012). Srgap37“ mice present a neurodevelopmental disorder with schizophrenia-related intermediate phenotypes. FASEB J. 26, 4418–28.

Wang, K., Cheung, E., Gong, Q.-Y., and Chan, R. (2011). Semantic processing disturbance in patients with schizophrenia: a meta-analysis of the N400 component. PLoS ONE 6, e25435. doi:10.1371/journal.pone.0025435.

Wang, R. (2011). Dissecting the Genetic Basis of Convergent Complex Traits Based on Molecular Homoplasy. PhD, Duke University.

Wang, Y., Huang, Y., Peng, M., Cong, Z., Li, X., Lin, A., et al. (2015). Association between Silent Information Regulator 1 (SIRT1) gene polymorphisms and schizophrenia in a Chinese Han population. Psychiatry Res. 225, 744–5.

Weiss, A.P., Dewitt, I., Goff, D., Ditman, T., and Heckers, S. (2005). Anterior and posterior hippocampal volumes in schizophrenia. Schizophr. Res. 73, 103–112. doi: 10.1016/j.schres.2004.05.018.

Weisz, N., Wuhle, A., Monittola, G., Demarchi, G., Frey, J., Popov, T., et al. (2014). Prestimulus oscillatory power and connectivity patterns predispose conscious somatosensory perception. Proc. Natl. Acad. Sci. U.S.A. 111, E417–e425.

Willemsen, M. H., Nijhof, B., Fenckova, M., Nillesen, W. M., Bongers, E. M., Castells-Nobau A, et al. (2013). GATAD2B loss-of-function mutations cause a recognisable syndrome with intellectual disability and are associated with learning deficits and synaptic undergrowth in Drosophila. J. Med. Genet. 50, 507–14.

Williams, L.E., Light, G.A., Braff, D.L., Ramachandran, V.S. (2010). Reduced multisensory integration in patients with schizophrenia on a target detection task. Neuropsychologia 48, 3128–3136.

Williams, N. A., Close, J. P., Giouzeli, M., and Crow, T. J. (2006). Accelerated evolution of Protocadherin11X/Y: A candidate gene-pair for cerebral asymmetry and language. Am. J. Med. Genet. B Neuropsychiatr. Genet. 141, 623–33.

Williams, S., and Boksa, P. (2010). Gamma oscillations and schizophrenia. J. Psychiatry Neurosci. 35, 75–77.

Wilson, N. K., Lee, Y., Long, R., Hermetz, K., Rudd, M. K., Miller, R., et al. (2011). A novel microduplication in the neurodevelopmental gene SRGAP3 that segregates with psychotic illness in the family of a COS proband. Case Rep. Genet. 2011, 585893.

Wilson, N. K., Lee, Y., Long, R., Hermetz, K., Rudd, M. K., Miller, R., et al. (2011). A novel microduplication in the neurodevelopmental gene SRGAP3 that segregates with psychotic illness in the family of a COS proband. Case Rep Genet. 2011, 585893.

Wisniewska, M. B. (2013). Physiological role of β-catenin/TCF signaling in neurons of the adult brain. Neurochem. Res. 38, 1144–55.

Wockner, L. F., Noble, E. P., Lawford, B. R., Young, R. M., Morris, C. P., Whitehall, V. L., et al. (2014). Genome-wide DNA methylation analysis of human brain tissue from schizophrenia patients. Transl. Psychiatry 4, e339.

Worthey, E. A., Raca, G., Laffin, J. J., Wilk, B. M., Harris, J. M., Jakielski, K. J., et al. (2013) Whole-exome sequencing supports genetic heterogeneity in childhood apraxia of speech. J. Neurodev. Disord. 5, 29.

Xu, T., Stephane, M., Parhi, K.K. (2012). Selection of abnormal neural oscillation patterns associated with sentence-level language disorder in schizophrenia. Conf. Proc. IEEE Eng. Med. Biol. Soc. 2012, 4923–6. doi: 10.1109/EMBC.2012.6347098.

Xu, T., Stephane, M., Parhi, K.K. (2013). Multidimensional analysis of the abnormal neural oscillations associated with lexical processing in schizophrenia. Clin. EEG Neurosci. 44, 135–143.

Yamada, K., Iwayama, Y., Hattori, E., Iwamoto, K., Toyota, T., Ohnishi, T., et al. (2011). Genome-wide association study of schizophrenia in Japanese population. PLoS One 6, e20468.

Yamaguchi, Y., Lee, Y. A., and Goto, Y. (2015). Dopamine in socioecological and evolutionary perspectives: implications for psychiatric disorders. Front. Neurosci. 9, 219.

Yolken, R.H., Severance, E.G., Sabunciyan, S., Gressitt, K.L., Chen, O., Stallings, C., et al. (2015). Metagenomic sequencing indicates that the oropharyngeal phageome of individuals with schizophrenia differs from that of controls. Schizophr. Bull. 41, 1153–1161.

Zhang, B., Xu, Y. H., Wei, S. G., Zhang, H. B., Fu, D. K., Feng, Z. F., et al. (2014). Association study identifying a new susceptibility gene (AUTS2) for schizophrenia. Int. J. Mol. Sci. 15, 19406–16.

Zhang, J., Liao, G., Liu, C., Sun, L., Liu, Y., Wang, Y., et al. (2011). The association of CLOCK gene T3111C polymorphism and hPER3 gene 54-nucleotide repeat polymorphism with Chinese Han people schizophrenics. Mol. Biol Rep. 38, 349–54.

Zweier, C. (2012). Severe intellectual disability associated with recessive defects in CNTNAP2 and NRXN1. Mol. Syndromol. 2, 181–185.

